# The role of disulfide bonds in the GluN1 subunit in the early trafficking and functional properties of GluN1/GluN2 and GluN1/GluN3 NMDA receptors

**DOI:** 10.1101/2025.10.07.680896

**Authors:** Jakub Netolicky, Seungha Lee, Petra Zahumenska, Marharyta Kolcheva, Anna Misiachna, Kristyna Rehakova, Stepan Kortus, Jae-man Song, Katarina Hemelikova, Emily Langore, Jovana Doderović, Marek Ladislav, Young Ho Suh, Martin Horak

## Abstract

*N*-methyl-D-aspartate receptors (NMDARs) are ionotropic glutamate receptors essential for excitatory neurotransmission. Previous studies proposed the existence of four disulfide bonds in the GluN1 subunit; however, their role in NMDAR trafficking remains unclear. Our study first confirmed the existence of four disulfide bonds in the GluN1 subunit using biochemistry in human embryonic kidney 293T (HEK293T) cells. Disrupting the individual disulfide bonds by serine replacements produced the following surface expression trend for GluN1/GluN2A, GluN1/GluN2B, and GluN1/GluN3A receptors: wild-type (WT) > GluN1-C744S-C798S > GluN1-C79S-C308S > GluN1-C420S-C454S > GluN1-C436S-C455S subunits. Electrophysiology revealed altered functional properties of NMDARs with disrupted disulfide bonds, specifically an increased probability of opening (Po) at the GluN1-C744S-C798S/GluN2 receptors. Synchronized release from the endoplasmic reticulum confirmed that disruption of disulfide bonds impaired early trafficking of NMDARs in HEK293T cells and primary hippocampal neurons prepared from Wistar rats of both sexes (embryonic day 18). The pathogenic GluN1-C744Y variant, associated with neurodevelopmental disorder and seizures, caused reduced surface expression and increased Po at GluN1/GluN2 receptors, consistent with findings for the GluN1-C744S-C798S subunit. The FDA-approved memantine inhibited GluN1-C744Y/GluN2 receptors more potently and with distinct kinetics compared to WT GluN1/GluN2 receptors. We also observed enhanced NMDA-induced excitotoxicity in hippocampal neurons expressing the GluN1-C744Y subunit, which memantine reduced more effectively compared to the WT GluN1 subunit. Lastly, we demonstrated that the presence of the hGluN1-1a-C744Y subunit counteracted the effect of the hGluN3A subunit on decreasing dendritic spine maturation, consistent with the reduced surface delivery of the NMDARs carrying this variant.

**Significance statement:** Our findings highlight the critical role of disulfide bonds in the GluN1 subunit in regulating trafficking and function of major conventional (GluN1/GluN2A, GluN1/GluN2B) and unconventional (GluN1/GluN3A) diheteromeric NMDAR subtypes in the postnatal forebrain. We further demonstrated that the pathogenic GluN1-C744Y variant reduces surface expression of all studied NMDARs, as well as increases the probability of opening (Po) of the GluN1/GluN2 receptors, leading to heightened NMDA-induced excitotoxicity in hippocampal neurons. Additionally, we introduced an ARIAD-based system for the synchronized release of NMDARs from the endoplasmic reticulum in hippocampal neurons. This system provides a powerful tool for studying pathogenic GRIN gene variants and addresses the current lack of molecular methods for analyzing the early trafficking of NMDARs.

## Introduction

*N*-methyl-D-aspartate receptors (NMDARs) mediate excitatory neurotransmission and synaptic plasticity in the mammalian central nervous system (CNS). They are formed as heterotetramers composed of two obligatory GluN1 subunits together with two GluN2 (GluN2A-GluN2D) and/or GluN3 (GluN3A-GluN3B) subunits (Traynelis et al., 2010; Vieira et al., 2020). All GluN subunits contain four membrane domains (M1 - M4), an extracellular amino-terminal domain (ATD), the S1 segment of the ligand-binding domain (LBD), an extracellular loop between M3 and M4 domains containing the S2 segment of the LBD, and an intracellular C-terminal domain (CTD) (Traynelis et al., 2010; Paoletti et al., 2013). The conventional NMDAR subtypes, containing GluN1/GluN2 subunits, are activated upon simultaneous binding of agonists, such as L-glutamate to the LBD of the GluN2 subunit, with co-agonists, such as glycine to the LBD of GluN1 subunit (Paoletti et al., 2013; Vieira et al., 2020). The unconventional NMDAR subtype, composed of GluN1/GluN3 subunits, is activated by the interaction of agonists such as glycine at the LBD of the GluN3 subunit, with desensitization mediated by glycine binding at the GluN1 subunit (Pérez-Otaño et al., 2016; Crawley et al., 2022).

Biogenesis of NMDARs begins with the translation of the GluN subunits in the rough endoplasmic reticulum (ER), where the unassembled GluN1-1a, GluN2, and GluN3 subunits are retained due to the presence of ER retention signals (Okabe et al., 1999; Standley et al., 2000; Pérez-Otaño et al., 2001; Scott et al., 2001; Matsuda et al., 2003; Hawkins et al., 2004; Horak et al., 2008; Horak and Wenthold, 2009). Previous studies proposed the presence of various disulfide bonds within the GluN subunits likely formed in the ER, including four disulfide bonds (C79-C308, C420-C454, C436-C455, C744-C798) in the GluN1 subunit (Laube et al., 1993; Choi et al., 2001; Lipton et al., 2002; Furukawa and Gouaux, 2003; Papadakis et al., 2004; Kaye et al., 2007). Interestingly, substitutions of the GluN1-C79 and GluN1-C308 residues reduced the surface expression of GluN1/GluN2B receptors in HEK293 cells by ∼50% (Papadakis et al., 2004) while increasing the EC_50_ value for NMDA by ∼25% (measured at GluN1/GluN2A receptors) (Choi et al., 2001). The disruption of the GluN1-C744-C798 disulfide bond by double serine replacement increased the current amplitudes of the GluN1/GluN3A receptors expressed in the HEK293 cells (Grand et al., 2018). Thus, previous studies indicated that the disruption of several disulfide bonds within the GluN1 subunit alters the functional properties of the NMDARs and may be one of the critical factors sensed by the ER quality control machinery during their biogenesis. However, this hypothesis has not been directly validated by comprehensively testing a complete series of mutated NMDARs with individually disrupted disulfide bonds.

Using a combination of biochemistry, microscopy, and electrophysiology in HEK293T cells and hippocampal neurons, we showed that the disruption of four known disulfide bonds within the GluN1 subunit resulted in the different degrees of the reduction of surface expression of mutated NMDARs, likely on the ER level. Our results also showed that the pathogenic GluN1-C744Y variant, which disrupts the formation of the C744-C798 disulfide bond, reduced the surface expression of NMDARs and enhanced NMDA-induced excitotoxicity, likely by increasing the probability of opening (Po) of GluN1-C744Y/GluN2 receptors. Given the predominantly gain-of-function effect associated with this variant, which is likely the cause of neurodevelopmental disorder and seizures, we tested an FDA-approved memantine as a potential open-channel blocker. Memantine demonstrated greater potency and altered inhibitory kinetics at GluN1-C744Y-containing NMDARs, suggesting its potential as a targeted treatment option.

## Material and Methods

### Molecular biology

We used DNA expression vectors containing rat versions of yellow fluorescent protein (YFP)-GluN1-1a (NP_058706.1), GluN1-4a (NP_001257539.1), GluN2A or green fluorescent protein (GFP)-GluN2A (NP_036705), GluN2B or GFP-GluN2B (NP_036706), and GFP-GluN3A (NP_612555.1) subunits. Human versions of YFP-GluN1-1a (YFP-hGluN1-1a; NP_015566.1), GluN1-4a (hGluN1-4a; NP_000823.4), GluN2A (hGluN2A; NP_001127879.1), GluN2B (hGluN2B; NP_000825.2), and GluN3A (hGluN3A; NP_597702) subunits were reported previously (Kaniakova et al., 2012; Skrenkova et al., 2019, 2020). The ARIAD-mNeonGreen-GluN1-1a (ARIAD-GluN1-1a) construct containing the ARIAD sequence (Hangen et al., 2018) and a rat GluN1-1a subunit with a mNEONGreen sequence inserted after the 21st amino acid residue was subcloned into the pcDNA3 expression vector. The ARIAD-mNeonGreen-hGluN1-1a (ARIAD-hGluN1-1a) construct was generated by introducing N159S, R212K, I267L, and M415L mutations in the GluN1-1a subunit (Skrenkova et al., 2020). The construction of the lentiviral FHUGW vector containing the wild-type (WT) YFP-hGluN1-1a subunit has also been described previously (Skrenkova et al., 2020). The amino acid substitutions were performed using the QuikChange site-directed mutagenesis kit (Agilent Technologies) and verified by sequencing the entire coding regions of the GluN1 subunits.

### HEK293T cells, lentiviruses, and primary hippocampal neurons

HEK293T cells (RRID: CVCL_0063, LGC Standards Sp. z.o.o.; ATCC® CRL-3216™; until passage number 10) were cultured in Opti-MEM I medium with 5% fetal bovine serum (FBS; both from Thermo Fischer Scientific). For electrophysiology, HEK293T cells grown in 24-well plates were transfected with 50 μl of Opti-MEM I medium containing 0.9 μg of DNA vectors encoding GFP (pQBI25; Takara), GluN1, GluN2 or GluN3A subunits (in a 1:1:1 ratio), and 0.9 μl of PolyMag reagent (OZBiosciences). After 20 min on a magnetic plate, HEK293T cells were trypsinized and resuspended in Opti-MEM I medium with 1% FBS, 2 mM MgCl_2_, and 3 mM kynurenic acid (to reduce excitotoxicity). For microscopy, HEK293T cells cultured on coverslips in 12-well plates were transfected with 50 μl of Opti-MEM I medium containing 0.45 μg of DNA vectors encoding GluN1, GluN2 or GluN3A subunits (at a ratio of 1: 2 for the YFP- or GFP-GluN subunit versus the untagged GluN subunit), and 1 μl of Lipofectamine 2000 reagent (Thermo Fischer Scientific). Similarly, DNA vectors containing ARIAD-(h)GluN1-1a constructs were transfected at a 1:2 ratio with untagged (h)GluN2A or (h)GluN3A subunits. HEK293T cells were cultured in 6-well plates and transfected with 0.5 μg of WT or mutant GluN1-4a subunit using the jetOPTIMUS reagent (Polyplus) for biochemistry experiments.

Lentiviral particles were prepared in the HEK293T cells by co-transfecting FHUGW vectors encoding YFP-hGluN1-1a or YFP-hGluN1-1a-C744Y subunits along the DNA vectors containing envelope expressing VSV glycoprotein (pVSVG), and envelope plasmids Rev and GAG/Pol (pΔ8.9) using Lipofectamine 2000 reagent. Viral supernatants were collected 36-48 hr after the end of transfection, and lentiviral particles were concentrated using 100 kDa Amicon Ultra centrifugal concentrators (Millipore). Concentrated supernatants were frozen at -80°C.

All animal experiments were performed in accordance with the ARRIVE guidelines and the European Commission Council Directive 2010/63/EU for animal testing. Primary cultures of hippocampal neurons were prepared from rats of both sexes of the Sprague-Dawley strain at embryonic day 18. Hippocampi were prepared in a chilled dissection medium containing Hanks’ balanced salt solution supplemented with 10 mM HEPES (pH 7.4), followed by incubation in a dissection medium supplemented with 0.05% trypsin and 0.1 mg/ml DNase I (Merck) for 20 min at 37°C. Dissociated cells were plated on poly-L-lysine-coated glass coverslips at a density of 2 × 10,000 cells per cm^2^ in a medium containing minimal essential medium (MEM) supplemented with 10% heat-inactivated horse serum, N2 supplement (1x), 1 mM sodium pyruvate, 20 mM D-glucose, 25 mM HEPES, and 1% penicillin-streptomycin (all from Thermo Fischer Scientific). After 3 hr, the plating medium was replaced with Neurobasal medium with 2% B-27 and 2 mM L-glutamine (all from Thermo Fischer Scientific), which was exchanged (∼50%) with fresh medium every 3-4 days (Lichnerova et al., 2015). Hippocampal neurons were transfected with DNA vectors containing genes for GluN subunits using Lipofectamine 2000 on *in vitro* day (DIV) 14 or infected using lentiviral particles on DIV4.

### Biochemistry

To evaluate the presence of disulfide bonds, HEK293T cells transfected with WT or mutant GluN1-4a subunits were harvested in TE buffer (50 mM Tris-HCl, pH 7.5, 1 mM EDTA) containing protease inhibitor cocktails (GenDEPOT) and sonicated. After solubilization with 1% SDS, the lysates were centrifuged at 20,000 × *g* for 15 min at 4J to remove insoluble materials. The supernatants were sequentially treated with 50 mM N-ethylmaleimide (NEM), 50 mM dithiothreitol (DTT), and/or 5 mM methoxypolyethylene glycol maleimide (PEG-mal). Each reaction was conducted for 1 h at room temperature. After the NEM or DTT steps, proteins were precipitated with chloroform/methanol/water (50:12.5:37.5, v/v/v) treatment to remove residual NEM or DTT from the lysates. The protein pellet was resuspended with TE buffer containing 1% SDS (Chen et al., 2016). The reaction was terminated by adding 6 × Laemmli buffer, and the migration of GluN1 subunits was analyzed by western blotting using an anti-GluN1 antibody (mouse anti-GluN1-NT, 1:10000; Merck).

For co-immunoprecipitation, primary hippocampal neurons were infected with lentiviruses expressing WT YFP-hGluN1-1a or YFP-hGluN1-1a-C744Y subunits for 48 h. Neurons were then harvested in TE buffer (50 mM Tris-HCl, pH 8.8; 2 mM EDTA) containing protease inhibitor cocktails, sonicated, and solubilized with 0.5% sodium deoxycholate. After neutralization with TE buffer at pH 6.8, the lysates were centrifuged at 20,000 g for 15 min at 4J. The supernatants were incubated with 0.5 mg of anti-GFP antibody (Invitrogen, Cat# A11122; RRID: AB_221569) for 1 h at 4J. The resulting immune complexes were further incubated with 20 mL of protein A-Sepharose beads (Sigma-Aldrich, Cat# P3391) for 3 h at 4J. After three washes, bound proteins were eluted in 6x Laemmli buffer and analyzed by western blotting using the following antibodies: anti-GluN2A (Millipore, Cat# 05-901R; RRID: AB_11215116), anti-GluN2B (Sigma-Aldrich, Cat# M265; RRID: AB_260487), anti-GluN3A (Millipore, Cat# 07-356; RRID: AB_2112620), and anti-GFP (Invitrogen, Cat# A11122; RRID: AB_221569).

### Microscopy

Immunofluorescence labeling in HEK293T cells was performed 24 hours after completion of transfection; hippocampal neurons were labeled 24 hours after transfection (DIV15) or 10 days after the lentiviral infection (DIV14) (Skrenkova et al., 2020; Kolcheva et al., 2023). Primary and secondary antibodies were diluted in a blocking solution of PBS and 0.2% bovine serum albumin (BSA; w/v; Merck). Cells were washed with ice-cold PBS and incubated with primary antibodies (rabbit anti-GFP, 1:1000; Millipore; Cat# AB3080P; RRID: AB_2630379) for 15 min on ice to label surface antigens, washed with a blocking solution, and incubated with a secondary antibody (goat anti-rabbit IgG conjugated with Alexa Fluor 555, 1:1000; Thermo Fisher Scientific; Cat# A-21428; RRID: AB_253584). Cells were washed with ice-cold PBS and fixed in a solution containing 4% paraformaldehyde (PFA; w/v; Merck) with 4% sucrose (w/v; Merck) in PBS for 15 min. Subsequently, the cells were permeabilized with 0.25% Triton X-100 (v/v; Merck) in PBS for 5 min and incubated with primary antibody (mouse anti-GFP, 1:1000; UC Davis/NIH NeuroMab Facility; Cat# N86/8; RRID: AB_2313651) and then secondary antibody (goat anti-mouse IgG conjugated with Alexa Fluor 488, 1:1000; Thermo Fisher Scientific; Cat# A-11001; RRID: AB_2534069) to label intracellular antigens (Skrenkova et al., 2020; Kolcheva et al., 2023). In experiments with ARIAD-GluN1-1a and ARIAD-hGluN1-1a constructs, cells were first incubated with ARIAD ligand at a concentration of 1 µM (AL; D/D Solubilizer; Takara) for the indicated times, fixed with PFA, labeled using primary (mouse anti-mNEON, 1:500; ChromoTek; Cat# 32f6-100; RRID: AB_2827566) and secondary (goat anti-mouse IgG conjugated with Alexa Fluor 555, 1:1000; Thermo Fisher Scientific; Cat# A-21422; RRID: AB_2535844) antibodies and then fixed in PFA; intracellular mNEON epitopes were not labeled (Netolicky et al., 2025). In co-localization experiments, the fixed and permeabilized cells labeled with primary (rabbit anti-GM130, 1:1000; Sigma-Aldrich Cat# G7295, RRID: AB_532244) and secondary (goat anti-rabbit IgG conjugated with Alexa Fluor 555, 1:1000; Thermo Fisher Scientific; Cat# A-21428; RRID: AB_253584) antibodies were briefly (5 min) fixed in PFA. The labeled cells were mounted using ProLong Antifade reagent (Thermo Fisher Scientific) (Skrenkova et al., 2019). Images were captured using an Olympus FV10i microscope with a 60×/1.35 oil immersion objective and analyzed using ImageJ 1.52N software (RRID: SCR_003070, National Institutes of Health). Total and surface fluorescence intensity was analyzed on the entire area of the transfected HEK293T cells. In the case of hippocampal neurons, 5 separate 10 µm segments of secondary or tertiary dendrites from a single neuron (total from n ≥ 4 neurons) were analyzed (Skrenkova et al., 2020; Kolcheva et al., 2023). The degree of co-localization of mNEON and GM130 signals was determined as the ratio between the intensity of the mNEON signal co-localizing with the GM130 signal and the intensity of the mNEON signal outside the co-localization region as described.

Twelve hours after transfection, HEK293T cells were treated with 1 mM L-glutamate and 100 μM glycine for 24 hours. Cleaved caspase-3 was detected by immunostaining using primary (rabbit anti-cleaved caspase-3, 1:250; Cell Signaling Technology; Cat# 9661; RRID: AB_2341188) and secondary (goat anti-rabbit IgG conjugated with Alexa Fluor 568, 1:500; Thermo Fisher Scientific; Cat# A-11036; RRID: AB_10563566) antibodies. Confocal images were acquired using a Zeiss LSM 800 microscope (RRID: SCR_015963). Image processing and fluorescence quantification were performed using Zen software (Zeiss) and ImageJ (NIH). Cleaved caspase-3-positive cells were manually counted in transfected HEK293T cells from randomly selected visual fields for each condition.

To examine the morphology of dendritic spines, primary hippocampal neurons (DIV 16) were fixed in PFA solution 48 hours after transfection with cytoplasmic mCherry, WT YFP-hGluN1-1a, or YFP-hGluN1-1a-C744Y and hGluN3A subunits. To visualize and classify dendritic spines, the mCherry signal was labeled with a primary rat anti-mCherry monoclonal antibody (1:500, Invitrogen, Cat# M11217) and then with a secondary antibody (goat anti-rat IgG conjugated with Alexa Fluor 555, 1:1000, Thermo Fisher Scientific, Cat# A-21434). The YFP tag was enhanced by primary rabbit anti-GFP antibody and goat anti-rabbit IgG conjugated with Alexa Fluor 488 (1:1000; Thermo Fisher Scientific; Cat# A-11008). Images were captured using an Olympus SpinSR10 microscope with a 60×/1.42 oil-immersion objective in super-resolution mode. Fifteen dendrites from at least ten neurons were analyzed for each condition. Spines were classified into one of the four morphological subtypes: mushroom, stubby, thin, and filopodia. Spines with a minimum head diameter of 0.35 μm and a head-to-neck ratio of at least 1.1 were classified as mushroom spines. Spines with a similar head diameter but lacking a distinct neck and directly attached to the dendrite were considered stubby. Spines with smaller, yet still detectable heads were categorized as thin, while spines lacking a visible head were classified as filopodia (Fiuza et al., 2013).

To examine the effect of synaptic activity on surface delivery of NMDARs, the cultured hippocampal neurons (DIV 14) were transfected with WT YFP-hGluN1-1a or YFP-hGluN1-1a-C744Y subunits together with either hGluN2A or hGluN2B subunits, and subsequently treated with bicuculline (20 μM; Hello Bio) or tetrodotoxin (TTX; 1 μM; Hello Bio) for 48 hours (Graves et al., 2021).

### Electrophysiology

The current responses elicited by applying the indicated concentrations of L-glutamate and/or glycine were recorded using the whole-cell patch-clamp technique ∼24-48 hours after transfection, using an amplifier Axopatch 200B (Molecular Devices) at holding potential -60 mV, using a rapid application system that achieves the solution exchange around the measured cell with a time constant of ∼15-20 ms. Electrophysiological recordings were filtered by an eight-pole Bessel filter capturing frequencies >2 kHz. Analog signals were digitized at 5 kHz using a Digidata 1550 A/D converter (Molecular Devices), and data were acquired using pCLAMP 10.7 software (RRID: SCR_011323; Molecular Devices). Borosilicate glass micropipettes with a tip resistance of 3-6 MΩ were prepared using a P-1000 puller (Sutter Instruments) and filled with an intracellular recording solution containing (in mM): 120 gluconic acid, 15 CsCl, 10 BAPTA, 10 HEPES, 3 MgCl_2_, 1 CaCl_2_ and 2 ATP-Mg salts (pH = 7.2 adjusted with CsOH). The extracellular recording solution (ECS) contained (in mM) 160 NaCl, 2.5 KCl, 10 HEPES, 10 D-glucose, 0.2 EDTA, and 0.7 CaCl_2_ (pH = 7.3 adjusted with NaOH). CGP-78608 (100 µM; Tocris) was diluted in dimethyl sulfoxide (DMSO). Dizocilpine (MK-801; 100 µM; Tocris) and memantine (100 µM; Hello Bio) were diluted in deionized water; all stock solutions were stored at -20°C. All electrophysiological measurements were performed at room temperature and recordings with a series resistance of <10 MΩ with 80% compensation were used in the analysis (Kaniakova et al., 2018; Kolcheva et al., 2023). For testing the redox sensitivity of NMDARs, we pre-applied for 2 min either the reducing agent dithiothreitol (DTT; 4 mM, Sigma-Aldrich) or the oxidizing agent 5,5 ′ -dithiobis-(2-nitrobenzoic acid) (DTNB; 0.5 mM, Sigma-Aldrich).

Concentration-dependent curves for L-glutamate and glycine were obtained using

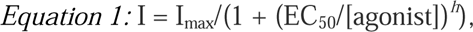

where I_max_ is the maximal steady-state current response elicited by agonist application, EC_50_ is the concentration of agonist eliciting half of the maximal response, [agonist] is the concentration of agonist, and h is the apparent Hill coefficient (Skrenkova et al., 2020; Kolcheva et al., 2023).

The time courses of inhibition by MK-801 (τ_w-MK-801_) and memantine (τ_on_ and τ_off_), the desensitization of the GluN1/GluN3A receptors (τ_w-des_) and of the L-glutamate-induced current responses (τ_on_ and τ_off_) were fitted with either a single-exponential or double-exponential function, and the corresponding weighted time constant (τ_w_, τ_on_ or τ_off_) was calculated using the following equation:

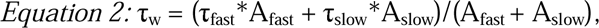

where τ_fast_ and A_fast_ correspond to the time constant and amplitude of the fast component, respectively, and τ_slow_ and A_slow_ correspond to the time constant and amplitude of the slow component, respectively.

The Po values were determined based on the inhibition kinetics by 1 µM MK-801 (Vyklicky et al., 2018). The onset of MK-801-induced inhibition was fitted to the following kinetic model without considering the ligand binding steps using Gepasi software (Mendes, 1993, 1997; Mendes and Kell, 1998):

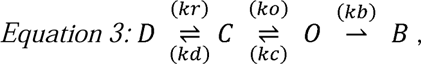

where D represents the desensitized state, C stands for the fully liganded close state, O is an open state, and B is the MK-801-induced blocked state. The determination of desensitization (D) alongside the kinetic constants for desensitization (kd) and resensitization (kr) were calculated using the following equations:

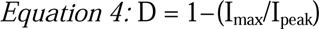

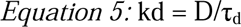

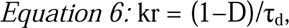

where I_peak_ represents the peak response, and I_max_ refers to the steady-state response. The single exponential function determined the onset of desensitization (τ_d_). The close rate (kc) was set at an arbitrary value of 250 s^−1^ (Kolcheva et al., 2023). The opening rate (ko) was set as a free parameter. As per established studies (Huettner and Bean, 1988; Jahr, 1992; Rosenmund et al., 1995), the MK-801 blocking rate (kb) was set at 25 μM^−1^ s^−1^. The binding of MK-801 was set as irreversible within the experimental timeframe. The calculation for microscopic Po was performed as follows:

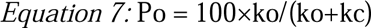

Concentration-dependent curves for memantine were obtained using

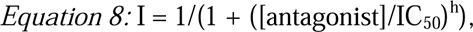

where IC_50_ is the concentration of memantine that caused half inhibition of the agonist-induced current, [antagonist] is the concentration of memantine, and *h* is the Hill coefficient.

### Excitotoxicity assay

Excitotoxicity was induced as previously described (Skrenkova et al., 2020). Briefly, cultured hippocampal neurons (DIV13) were incubated overnight in 10% MEM (Thermo Fisher Scientific) and 90% salt-glucose medium (SG) containing (in mM): 114 NaCl, 5.292 KCl, 1 MgCl_2_, 2 CaCl_2_, 10 HEPES, 30 D-glucose, and 0.5 sodium pyruvate, including 0.219% NaHCO_3_ and 0.1% phenol red. The following day, the medium was replaced with 100% SG medium, and the indicated concentrations of NMDA and glycine were added to the neurons. After 1 hour, this medium was replaced with 10% MEM and 90% SG medium. Twenty-three hours later, cell nuclei were labeled with Hoechst 33342 (5 µM; Cat# H1399; Molecular Probes) for 30 min. YFP-hGluN1 subunits in the fixed and permeabilized neurons were labeled with primary (rabbit anti-GFP; Millipore; Cat# AB3080P; RRID: AB_2630379) for) and secondary (Alexa Fluor 488-conjugated goat anti-rabbit Thermo Fisher Scientific; Cat# A-11008; RRID: AB_143165) antibodies. Images (1,024 × 1,024 pixels with a pixel size of 1.243 × 1.243 µm, covering a field of 1,272 × 1,272 µm) were acquired using an Olympus FV10i confocal microscope with a 60×/1.35 oil immersion objective; three separate images were acquired for each focused area: the YFP signal (to detect the infected cells), the Hoechst 33342 signal (to image single cell nuclei), and a wide-field image. The nuclear size was measured using ImageJ software (RRID: SCR_003070, v. 1.52p), and custom macro scripts were used to detect nuclei in infected cells automatically (identified by YFP expression). All measured nuclei sizes from a single experiment were plotted on a histogram covering control (cells with predominantly non-pyknotic nuclei) and individual test conditions (predominantly a mixture of cells with pyknotic and non-pyknotic nuclei). The histogram contained two distinguishable groups corresponding to cells with pyknotic and non-pyknotic nuclei. The Matlab (RRID: SCR_001622) function “fitgmdist()” was used to fit the histograms via two Gaussian functions. The data obtained under each condition were then processed using a Gaussian mixture model in the MatLab function “cluster()”, which allowed estimating the posterior probability of each measured nuclear region belonging to one of the distribution types and sorted the cells into two groups. The ratio of the number of cells in each group was then used to estimate the effect of mixing the two distributions and was expressed as the ratio of cells with pyknotic to non-pyknotic nuclei. The percentage of cells with pyknotic nuclei for each condition was then calculated as the ratio of cells classified as pyknotic to the total number of cells in each condition.

### Statistics

Statistical analysis of microscopic data was conducted on normalized and log-transformed data to stabilize variability against the mean. Outliers outside 1.5 times the interquartile range were removed, and normality was assessed using the D’Agostino-Pearson test. Statistical significance was determined using one-way or two-way ANOVA, followed by Tukey’s multiple comparison tests via Matlab (RRID: SCR_001622, Matlab 2022b; functions ‘anova1’, ‘anovan,’ and ‘multcompare’). Linear regression was analyzed using Matlab’s ‘robustfit’ function. Points with residuals ≥2 times the median absolute deviation (MAD) were identified as outliers. Regression quality was expressed by the coefficient of determination (R²) and 95% confidence intervals were calculated using the ‘predint’ function (’parameter functional’ set to ‘on’). Additionally, Pearson’s correlation coefficients (r) and corresponding p-values were determined using Matlab’s ‘corr’ function to validate the statistical significance of observed correlations. Both r and R² are shown to describe correlations. Summarized data were presented as mean ± standard error of the mean (SEM), and differences with a p-value <0.050 were considered statistically significant.

## Results

### The cysteine residues of the GluN1 subunit form four disulfide bonds with additional free thiol

Previous studies suggested the presence of four disulfide bonds within the GluN1 subunit, C79-C308, C420-C454, C436-C455, C744-C798 (Fig. 1*A,B*)(Laube et al., 1993; Lipton et al., 2002; Furukawa and Gouaux, 2003; Papadakis et al., 2004). We first aimed to verify the presence of these disulfide bonds by biochemical analysis in HEK293T cells transfected with WT and various cysteine-to-serine mutated GluN1-4a subunits. Our experimental strategy included the sequential treatment with NEM, DTT, and PEG-mal (Fig. 1*C*), followed by western blot analysis of the GluN1-4a subunit. NEM, a thiol-specific reagent, forms irreversible covalent thioether bonds with free sulfhydryls, while leaving the oxidized cystines (S-S) within the cysteine disulfide bonds intact. Thus, NEM blocks the reduced sulfhydryl groups (S-H) on cysteines. Due to its small molecular weight of 125 Da, NEM minimally affects protein mobility in polyacrylamide gels. Subsequent treatment with the reducing agent DTT reduces pre-formed disulfide bonds between cysteine residues through two sequential thiol-disulfide exchange reactions, exposing free thiols. Finally, PEG-mal covalently binds to these reduced sulfhydryls through thioether bonds. Although the molecular weight of the PEG moiety is 5 kDa, its addition increases the molecular weight of the target protein by ∼10-15 kDa (Chen et al., 2016). First, sequential treatment with NEM– DTT–PEG-mal revealed that the WT GluN1-4a subunit possesses both free cysteine residues and disulfide bonds between cysteine residues (Fig. 1*D*). GluN1-4a subunit treated with PEG-mal migrates more slowly (Fig. 1*D*, indicated by an asterisk) due to the attachment of PEG-mal to the free cysteine residues, confirming the presence of free thiols in GluN1-4a subunit (Fig. 1*D*, lane 1 vs. 2). GluN1-4a subunit treated with PEG-mal exhibited much slower migration following DTT treatment, because DTT reduces pre-existing disulfide bridges, exposing additional free thiols for PEG-mal attachment (Fig. 1*D*, lane 2 vs. 3, and lane 4 vs. 5). These results indicate the presence of multiple disulfide bonds in GluN1-4a subunit.

**Figure 1.**
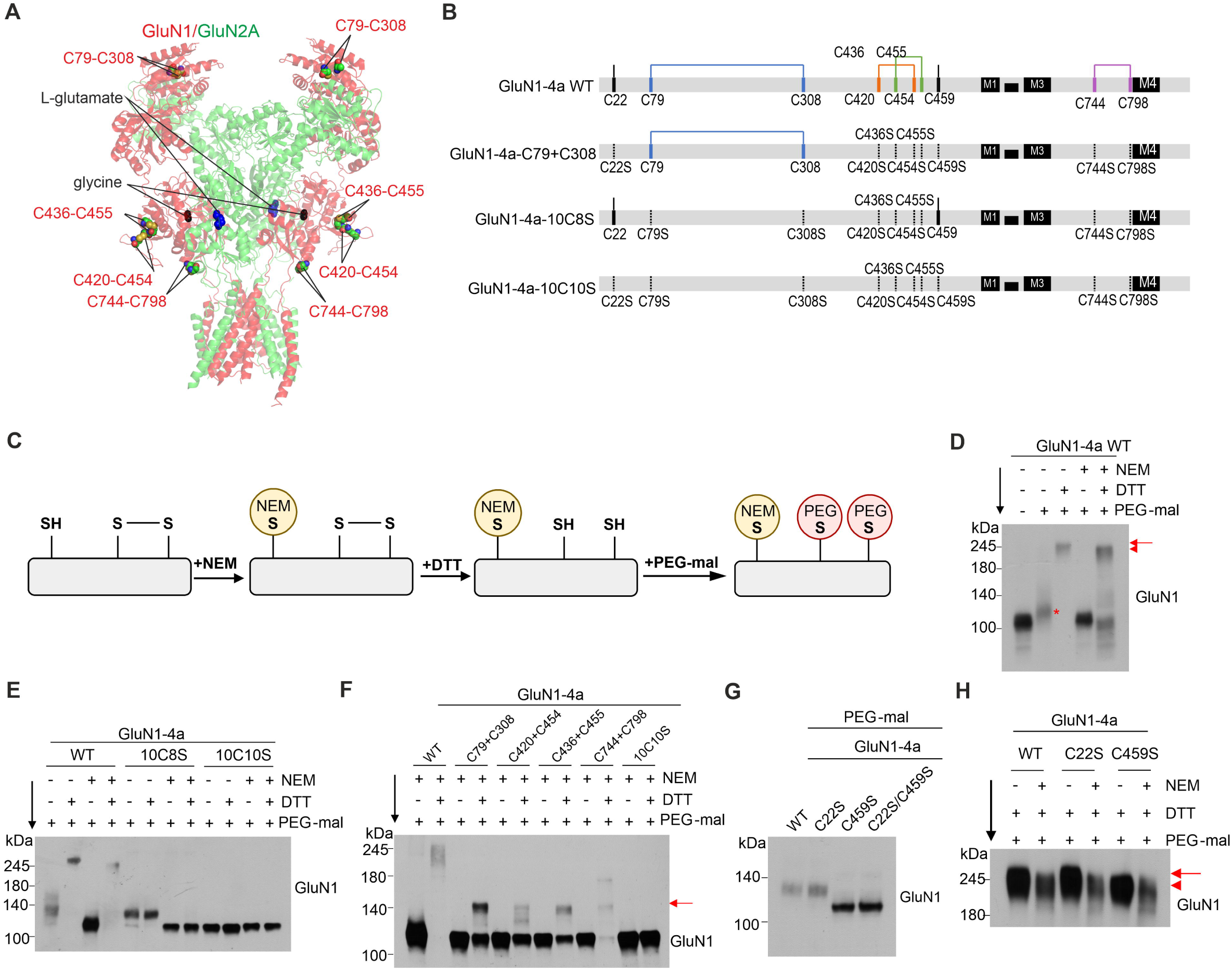
The cysteine residues of the GluN1 subunit form four disulfide bonds. ***A,*** The structural model of the NMDAR consists of GluN1 (in red) and GluN2A (in green) subunits (PDB:7EU7) shown with L-glutamate and glycine molecules bound within the LBDs. Cysteine residues potentially forming the disulfide bonds within the GluN1 subunit are shown as spheres and labeled in red. ***B,*** Schematic diagrams showing the positions of 10 cysteine (C) residues and four disulfide bonds in WT GluN1-4a subunit and cysteine-to-serine GluN1-4a subunit mutants that disrupt disulfide bonds. Each putative disulfide bond is represented by a solid line of the same color, while dashed lines indicate the positions of the cysteine-to-serine mutations. ***C,*** Schematic illustration of the biochemical strategy used to detect disulfide bonds in the GluN1-4a subunit. Following cell lysis, N-ethylmaleimide (NEM), dithiothreitol (DTT), and methoxypolyethylene glycol maleimide (PEG-mal) were applied sequentially. ***D,*** The gel migration pattern of the WT GluN1-4a subunit was analyzed by western blotting following sequential treatment with NEM–DTT–PEG-mal. The asterisk represents the slower migration of the GluN1 subunit due to PEG-mal attachment to free thiol residues. The red arrow and arrowhead indicate the slower and faster migration of the GluN1-4a subunit, respectively, depending on the masking of free thiols by NEM pre-treatment. ***E,*** Gel migration patterns of WT and mutant GluN1-4a subunits lacking disulfide bonds following sequential treatment with NEM–DTT–PEG-mal. ***F,*** Gel migration patterns of GluN1-4a C+C mutants, each containing only one disulfide bond between the designated cysteine pairs. The red arrow indicates the band shift from PEG-mal attachment to free thiols exposed by DTT treatment. ***G,*** Gel migration patterns of GluN1-4a subunit mutants lacking free thiols after PEG-mal treatment. ***H,*** Gel migration patterns of GluN1-4a subunit mutants lacking free thiols following sequential treatment with NEM–DTT–PEG-mal. The red arrow and arrowhead indicate the slower (lanes 1 and 3) and faster (lanes 2, 4, 5, and 6) migration of the GluN1-4a subunit, respectively. Full-length western blot images are provided in Figure S1.

Next, we generated the mutant GluN1-4a-10C8S subunit, which lacks all putative cysteine residues forming disulfide bonds, and the mutant GluN1-4a-10C10S subunit, which lacks all cysteine residues, by mutating cysteine to serine residues. The PEG-mal-treated GluN1-4a-10C8S subunit migrated at the same velocity as the PEG-mal-treated WT GluN1-4a subunit (Fig. 1*E*, lane 1 vs. 5), indicating that the putative free cysteine residues do not form disulfide bonds with the bridged cysteines. No band shift was observed following DTT treatment of the GluN1-4a-10C8S subunit (Fig. 1*E*, lane 5 vs. lane 6, and lane 7 vs. lane 8), indicating that no additional disulfide bonds are present in the GluN1-4a subunit beyond the maximum of four disulfide bonds. The band shift of PEG-mal-treated GluN1-4a-10C8S subunit compared to PEG-mal-treated GluN1-4a-10C10S subunit indicates the presence of free cysteine residues, such as C22 or C459 (Fig. 1*E*, lane 5 vs. 9). Furthermore, NEM pre-treatment did not alter the migration velocity of PEG-mal-treated GluN1-4a-10C10S subunit (Fig. 1*E*, lane 9 vs. lane 11), confirming the absence of additional free cysteine residues in the GluN1-4a-10C10S subunit. To identify the exact pairs of disulfide bonds in the GluN1-4a subunit, we generated individual C+C mutants, each containing one putative cysteine disulfide bond pair, with all other cysteine residues mutated to serine residues (Fig. 1*B*, e.g., C79+C308). After DTT–PEG-mal treatment, all four C+C mutants migrated more slowly compared to the DTT-untreated mutants (Fig. 1*F*), indicating that the GluN1-4a subunit contains four disulfide bonds, each corresponding to a pair in the C+C mutants, as illustrated in Fig. 1*B*.

Finally, we investigated the presence of free cysteine residues in the GluN1-4a subunit containing the GluN1-4a-C22S and/or GluN1-4a-C459S mutations through PEG-mal attachment. The GluN1-4a-C22S subunit exhibited the same migration pattern as the WT GluN1-4a subunit (Fig. 1*G*). In contrast, the GluN1-4a-C459S subunit migrated at the same velocity as the double mutant GluN1-4a-C22S/C459S subunit. This indicates that, unlike the GluN1-4a-C459 residue, the GluN1-4a-C22 residue does not harbor a free thiol group. These findings suggest that the GluN1-4a-C22 residue either forms an additional disulfide bond with another cysteine residue or is absent in the mature GluN1-4a subunit. When free thiols in GluN1-4a subunit were blocked by NEM pre-treatment, subsequent DTT–PEG-mal treatment resulted in faster migration of WT GluN1-4a or GluN1-4a-C22S subunits (Fig. 1*H*, lane 1 vs. 2, and lane 3 vs. 4), whereas it did not affect the migration velocity of the mutant GluN1-4a-C459S subunit (Fig. 1*H*, lane 5 vs. 6). These results suggest that the C22 residue is absent in the mature GluN1-4a subunit, rather than forming an additional disulfide bond.

### Disruption of disulfide bonds within the GluN1 subunit reduces surface expression and alters the functional properties of GluN1/GluN2 receptors

We next investigated whether disulfide bonds in the GluN1 subunit regulate the surface expression of GluN1/GluN2A receptors expressed in HEK293T cells. We created a series of double mutants containing C79S-C308S, C420S-C454S, C436S-C455S, and C744S-C798S substitutions in the YFP-GluN1-1a subunit (which is not transported to the cell surface, unless co-expressed with the GluN2 or GluN3 subunits) or in the untagged GluN1-4a subunit (which is transported to the cell surface even in the absence of the GluN2 and GluN3 subunits) (Standley et al., 2000; Scott et al., 2001). We co-expressed the WT and mutant YFP-GluN1-1a subunits with the untagged WT GluN2A subunit (Fig. 2*A*), and the WT and mutant GluN1-4a subunits together with the WT GFP-GluN2A subunit in HEK293T cells and determined their total and surface expression levels by immunofluorescent labeling with an anti-GFP antibody. Our microscopy analysis with both GluN1/GluN2A receptor combinations showed that the disruption of any of the four disulfide bonds in the GluN1 subunit reduced their surface expression, in the following order: WT GluN1/GluN2A > GluN1-C744S-C798S/GluN2A > GluN1-C79S-C308S/GluN2A > GluN1-C420S-C454S/GluN2A > GluN1-C436S-C455S/GluN2A (Fig. 2*B* and S2). These findings showed that both tags (e.i., YFP or GFP) and splice variants of the GluN1 subunit had no discernible impact on the microscopy results, as both GluN1/GluN2A receptor combinations practically mirrored each other.

**Figure 2.**
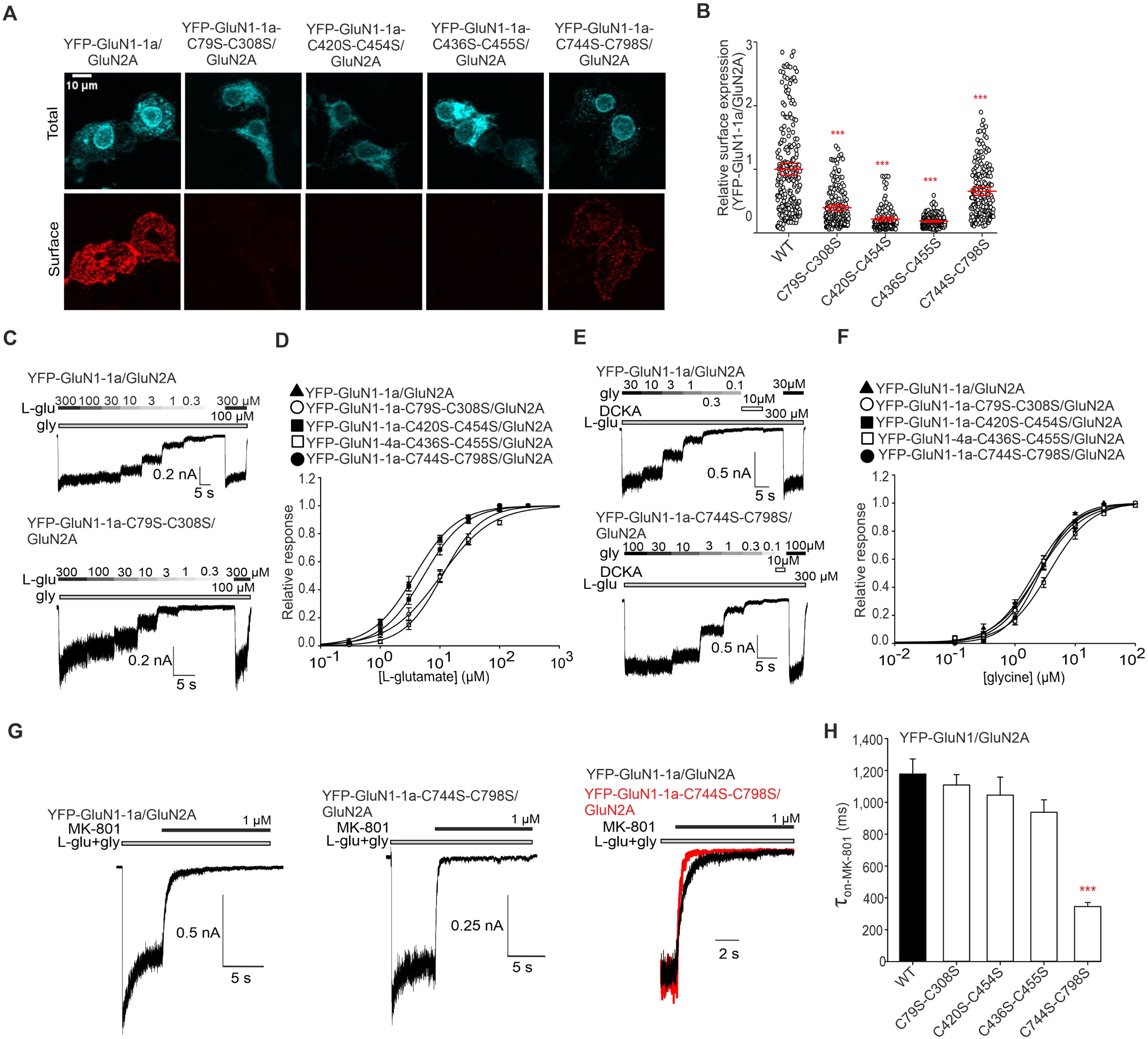
Mutations of cysteine residues forming disulfide bonds within the GluN1 subunit affect surface expression and functional properties of GluN1/GluN2A receptors. ***A,*** Representative images of HEK293T cells co-expressing the WT or mutated YFP-GluN1-1a subunit and the GluN2A subunit. The total and the cell surface number of YFP-GluN1-1a subunits (top and bottom row, respectively) were labeled using an anti-GFP antibody 24 hours after the transfection. ***B,*** Summary of the relative surface expression of NMDARs consisting of WT or mutated YFP-GluN1-1a subunit co-expressed with the GluN2A subunit, measured using fluorescence microscopy; one-way ANOVA, *F*(4, 883) = 164.5, *p* < 0.001; post hoc Tukeýs tests:\*\*\**p < 0.001* for WT vs. mutated NMDARs. Data points represent individual cells (*n* ≥ 129), and the red box plot represents mean ± SEM. For the relative surface expression of NMDARs containing the WT or mutated GluN1-4a and the WT GFP-GluN2A subunits, see Figure S2 ***C, E,*** Representative whole-cell voltage-clamp recordings showing the concentration-response relationship of agonists in HEK293T cells expressing the indicated NMDARs. Currents were elicited by L-glutamate (L-glu) at the indicated concentrations in the continuous presence of 100 µM glycine (gly) (C), or the current was elicited by glycine at the indicated concentrations in the continuous presence of 300 µM L-glutamate (E). ***D, F,*** Normalized concentration-response curves for L-glutamate (D) and glycine (F) measured from HEK293T cells expressing NMDARs containing WT or mutated YFP-GluN1-1a (YFP-GluN1-4a) subunit together with the GluN2A subunit. The data were fitted using *Equation 1* (see Methods); for a summary of fitting parameters, see Tables 1 and 2; for correlations with relative surface expression, see Figures S3 and S4, respectively. ***G,*** Representative whole-cell voltage-clamp recordings from HEK293T cells expressing indicated NMDARs, showing the onset of MK-801 inhibition used to estimate the channels’ open probability (Po). MK-801 (1 µM, black bar) was applied in the continuous presence of 1 mM L-glutamate (L-glu, grey bar) and 100 µM glycine (gly, grey bar). On the right, the onset of inhibition induced by 1 µM MK-801 for both receptors is overlaid and scaled to WT receptor response. ***K,*** Summary of the τ_on-MK-801_ values for NMDARs consisting of WT or mutated YFP-GluN1-1a subunits co-expressed with the GluN2A subunit obtained by fitting the data using *Equation 2* (see Methods, Table 3); one-way ANOVA, *F*(4, 24) = 14.83*, p* < 0.001; post hoc Tukey’s tests: \*\*\**p* < 0.001 for WT vs. mutated NMDARs. For the correlation analysis of relative surface expression and Po values (see Methods), see Figure S5.

**Table 1.**
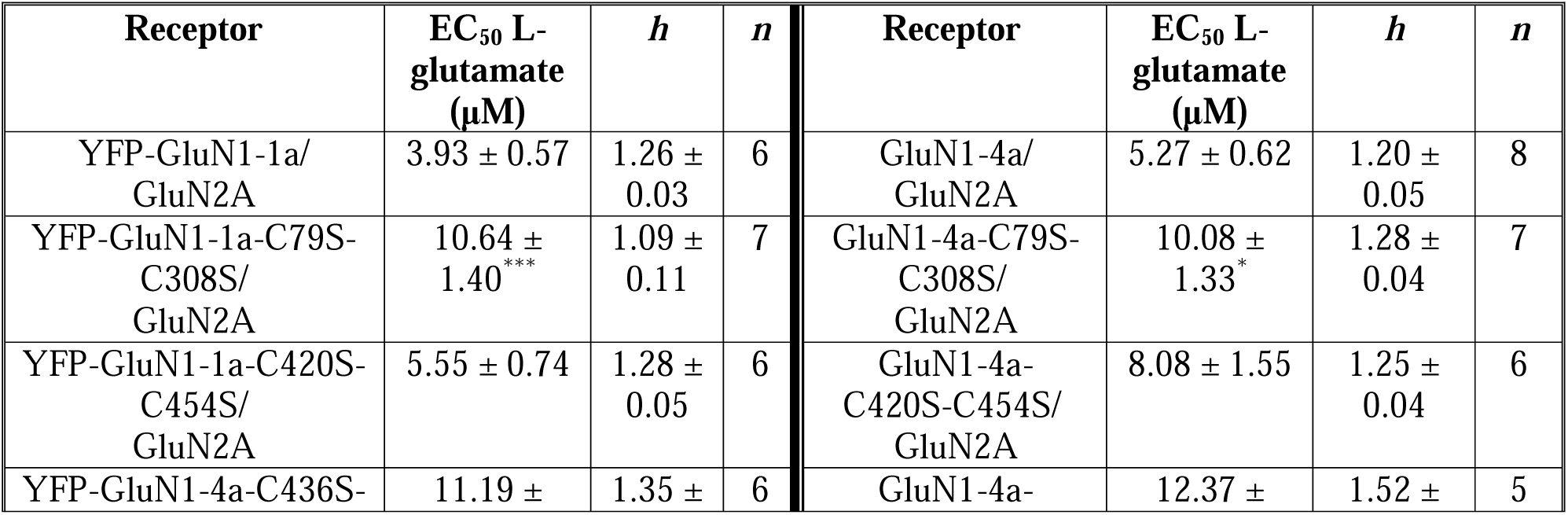
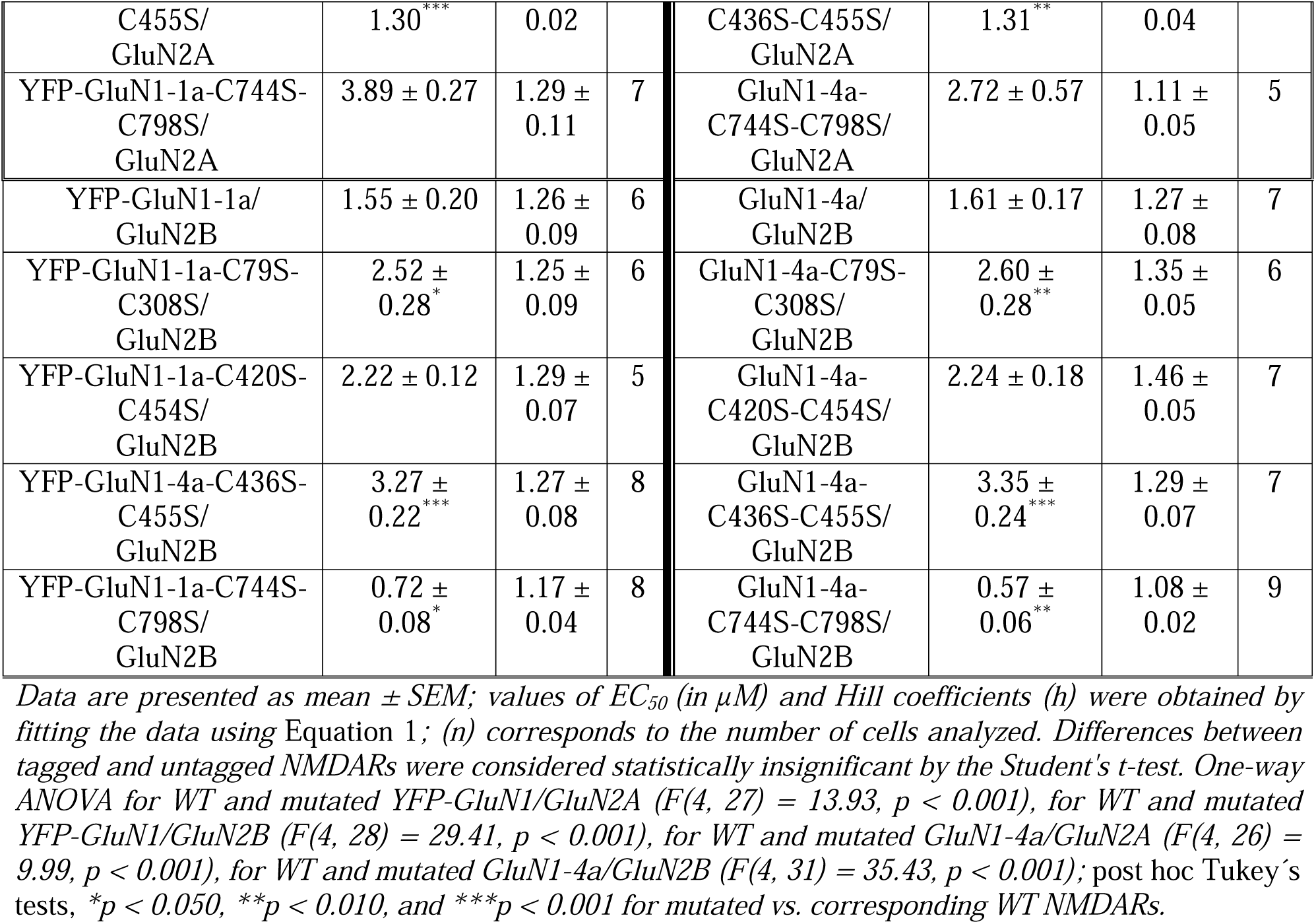
Summary of the L-glutamate concentration-response relationship analysis at WT and mutated recombinant NMDARs expressed in HEK293T cells with or without fluorescent tag (YFP or GFP).

**Table 2.**
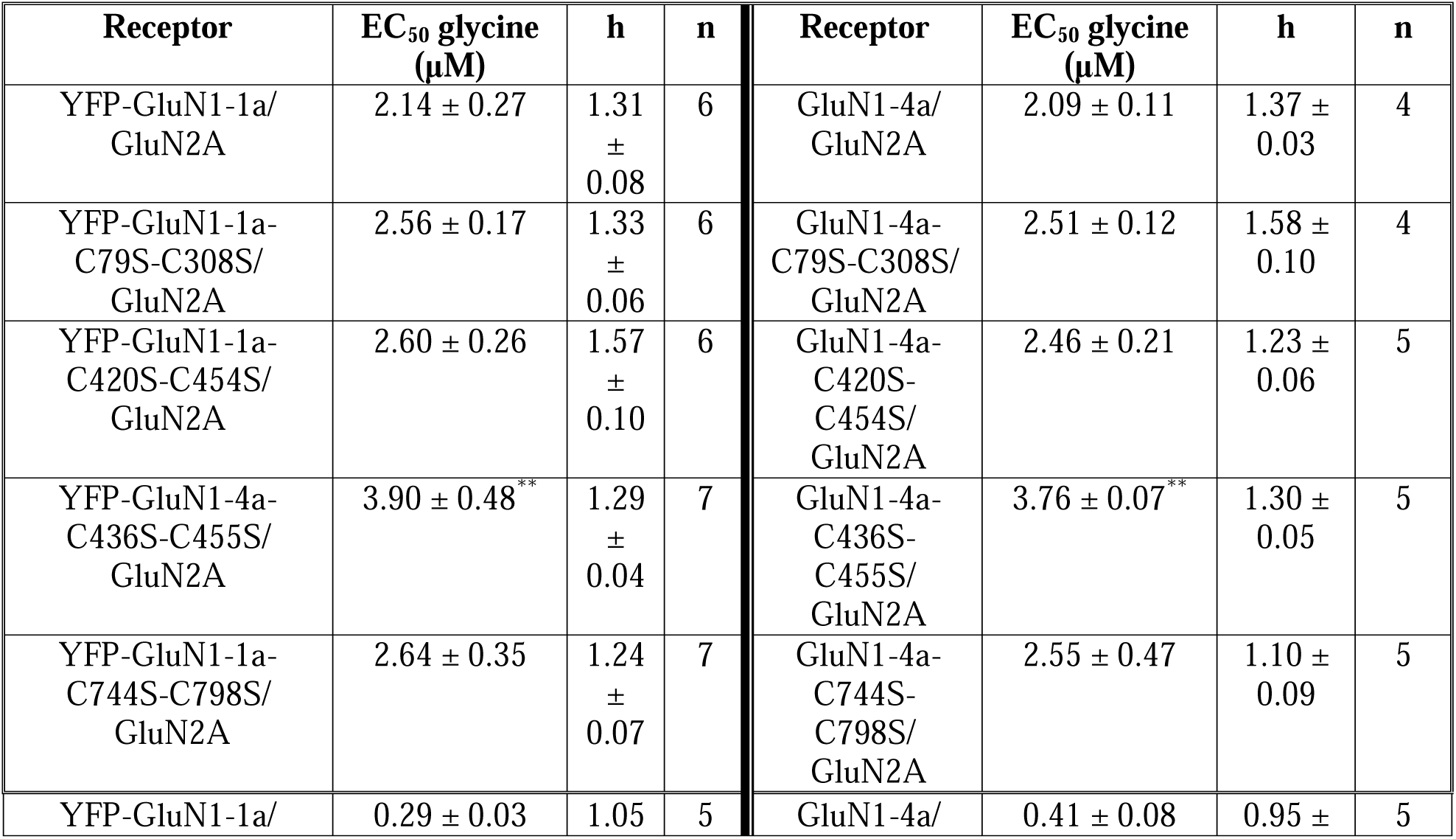
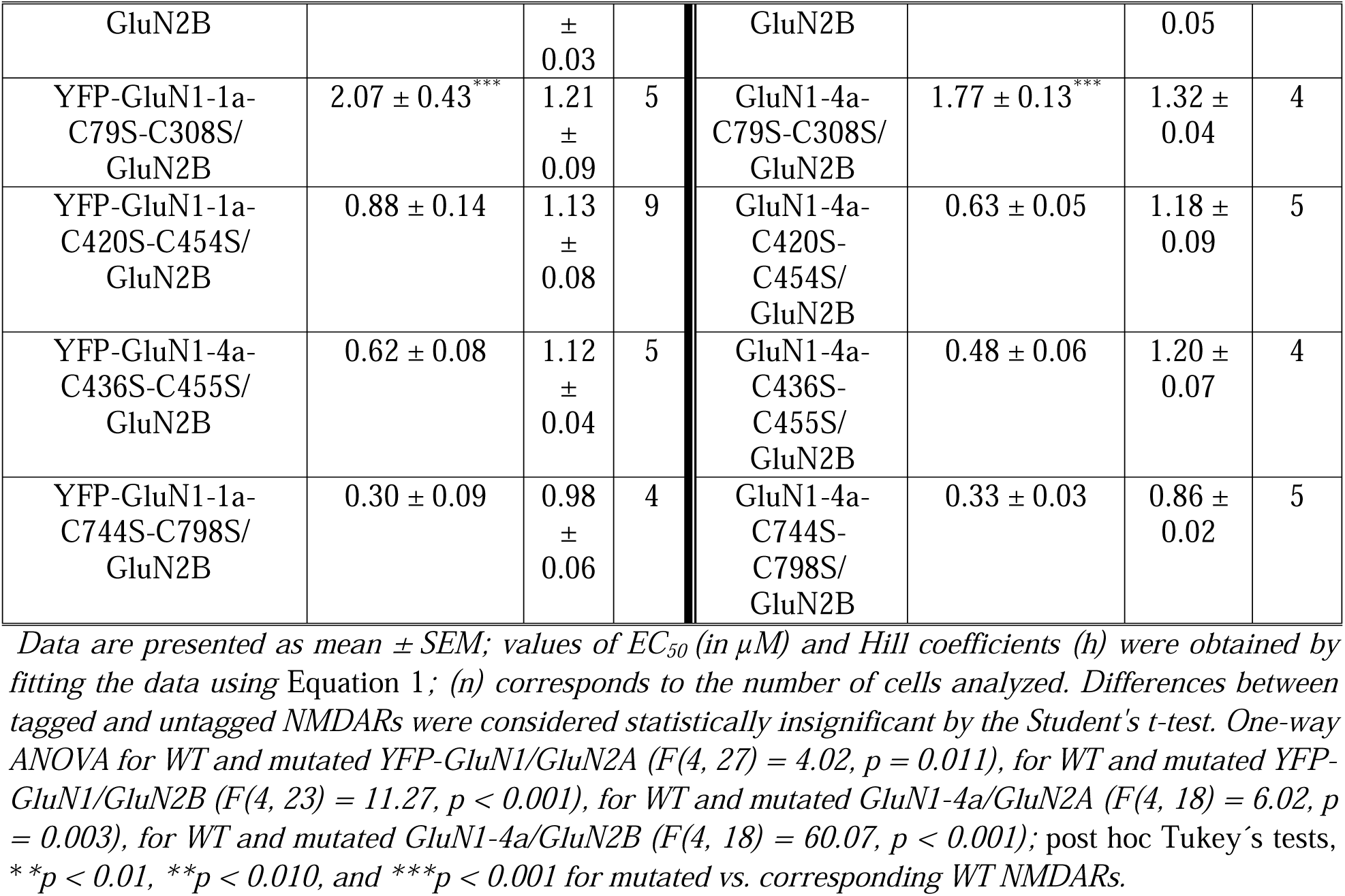
Summary of the glycine concentration-response relationship analysis at WT and mutated recombinant NMDARs expressed in HEK293T cells with or without fluorescent tag (YFP or GFP).

**Table 3.** Summary of the time constants and Po values estimated based on the inhibition onset of MK-801 at WT and mutated recombinant NMDARs expressed in HEK293T cells with or without fluorescent tag.

| Receptor | $\tau_{on-MK-801}$ (ms) | Po | $n$ | Receptor | $\tau_{on-MK-801}$ (ms) | $n$ |
| --- | --- | --- | --- | --- | --- | --- |
| YFP-GluN1-1a/<br>GluN2A | $1,178.37 \pm 93.82$ | $0.066 \pm 0.009$ | 6 | GluN1-4a/<br>GluN2A | $981.54 \pm 19.11$ | 5 |
| YFP-GluN1-1a-C79S-C308S/<br>GluN2A | $1,109.06 \pm 65.15$ | $0.055 \pm 0.010$ | 7 | GluN1-4a-C79S-C308S/<br>GluN2A | $1,038.07 \pm 140.54$ | 6 |
| YFP-GluN1-1a-C420S-C454S/<br>GluN2A | $1,045.30 \pm 112.94$ | $0.077 \pm 0.010$ | 6 | GluN1-4a-C420S-C454S/<br>GluN2A | $987.68 \pm 105.05$ | 6 |
| YFP-GluN1-4a-C436S-C455S/<br>GluN2A | $937.85 \pm 78.10$ | $0.077 \pm 0.007$ | 5 | GluN1-4a-C436S-C455S/<br>GluN2A | $910.61 \pm 85.39$ | 6 |
| YFP-GluN1-1a-C744S-C798S/<br>GluN2A | $345.28 \pm 25.68^{***}$ | $0.195 \pm 0.016^{***}$ | 5 | GluN1-4a-C744S-C798S/<br>GluN2A | $278.04 \pm 55.79^{**}$ | 5 |
| YFP-GluN1-1a/<br>GluN2B | $2,053.21 \pm 117.07$ | $0.024 \pm 0.002$ | 5 | GluN1-4a/<br>GluN2B | $2,307.47 \pm 151.45$ | 4 |
| YFP-GluN1-1a-C79S-C308S/<br>GluN2B | 3,010.09 ±<br>193.67** | 0.014 ±<br>0.001 | 5 | GluN1-4a-C79S-C308S/<br>GluN2B | 3,355.73<br>± 243.72* | 4 |
| YFP-GluN1-1a-C420S-C454S/<br>GluN2B | 2,612.49 ±<br>167.91 | 0.017 ±<br>0.001 | 6 | GluN1-4a-C420S-C454S/<br>GluN2B | 2,213.11 ±<br>189.99 | 6 |
| YFP-GluN1-4a-C436S-C455S/<br>GluN2B | 2,040.87 ±<br>282.26 | 0.026 ±<br>0.003 | 4 | GluN1-4a-C436S-C455S/<br>GluN2B | 1,988.70 ±<br>356.79 | 4 |
| YFP-GluN1-1a-C744S-C798S/<br>GluN2B | 765.01 ±<br>74.13*** | 0.055 ±<br>0.006*** | 6 | GluN1-4a-C744S-C798S/<br>GluN2B | 486.89 ±<br>100.90** | 4 |

Data are presented as mean ± SEM; values of $\tau_{w-MK-801}$ (in ms) were obtained by fitting the data using Equation 2, and values of $P_o$ were estimated based on fitting the inhibition onset of MK-801 to a kinetic model using Gepasi (see Methods); (n) corresponds to the number of cells analyzed. Differences between tagged and untagged NMDARs were considered statistically insignificant by Student's *t*-test. For $\tau_{on-MK-801}$ : one-way ANOVA for WT and mutated YFP-GluN1/GluN2A ( $F(4,24) = 14.83$ , $p < 0.001$ ), for WT and mutated YFP-GluN1/GluN2B ( $F(4,21) = 28.80$ , $p < 0.001$ ), for WT and mutated GluN1-4a/GluN2A ( $F(4,23) = 9.82$ , $p < 0.001$ ), for WT and mutated GluN1-4a/GluN2B ( $F(4,17) = 19.43$ , $p < 0.001$ ). For $P_o$ : one-way ANOVA for WT and mutated YFP-GluN1/GluN2A ( $F(4,24) = 26.32$ , $p < 0.001$ ), for WT and mutated YFP-GluN1/GluN2B ( $F(4,21) = 24.60$ , $p < 0.001$ ); post hoc Tukey's tests, \* $p < 0.05$ , \*\* $p < 0.010$ , and \*\*\* $p < 0.001$ for mutated vs. corresponding WT NMDARs.

We next asked whether GluN1/GluN2A receptors with disrupted disulfide bonds exhibit altered functional properties, including their sensitivity to L-glutamate, glycine, and Po. We employed whole-cell patch-clamp recordings from HEK293T cells transfected with WT and mutant YFP-GluN1-1a/GluN2A receptors; the current responses were elicited by rapid application of either L-glutamate at various concentrations in the presence of 100 µM glycine (Fig. 2*C*) or glycine at various concentrations in the presence of 300 µM L-glutamate (Fig. 2*E*). These electrophysiological measurements showed that HEK293T cells expressing WT and all mutant YFP-GluN1-1a/GluN2A receptors, except YFP-GluN1-1a-C436S-C455S/GluN2A receptor, produced sufficiently large current responses to construct concentration-dependent response curves and calculate EC_50_ values for L-glutamate (Fig. 2*D*) and glycine (Fig. 2*F*). To obtain sufficient current for the GluN1-C436S-C455S double mutant, we examined the YFP-GluN1-4a-C436S-C455S/GluN2A receptor instead. For control purposes, we performed electrophysiological measurements from HEK293T cells expressing untagged WT and mutant GluN1-4a/GluN2A receptors to determine whether the YFP tag or different GluN1 isoforms affected the EC_50_ values. These experiments showed that the YFP tag did not influence the EC_50_ values for L-glutamate (Table 1) or glycine (Table 2). Moreover, we observed that both combinations of i.) GluN1-C79S-C308S/GluN2A and GluN1-C436S-C455S/GluN2A receptors exhibited higher EC_50_ values for L-glutamate (Table 1), and ii.) GluN1-C436S-C455S/GluN2A receptors had higher EC_50_ values for glycine. Using linear regression, we observed no statistically significant correlation of surface expression to EC_50_ values for L-glutamate (r = -0.67, p = 0.21, R^2^ = 0.45, Fig. S3) and glycine (r = -0.69, p = 0.20, R^2^ = 0.47, Fig. S4) for WT and mutant YFP-GluN1/GluN2A receptors. Then, we employed an irreversible open-channel blocker of the NMDARs, MK-801, to estimate the Po values of WT and mutant GluN1/GluN2A receptors. We elicited current responses of WT and mutant YFP-GluN1-1a/GluN2A (Fig. 2*G*) and GluN1-4a/GluN2A receptors expressed in HEK293T cells using 1 mM L-glutamate in the presence of 100 µM glycine and then co-applied 1 µM MK-801. Our analysis showed that i.) both combinations of the GluN1-C744S-C798S/GluN2A receptors exhibited reduced onset time of MK-801 inhibition (τ_w-MK-801_) values compared to the corresponding WT GluN1/GluN2A receptors, and ii.) the presence of the YFP tag did not affect the τ_w-MK-801_ values measured for WT and mutant YFP-GluN1-1a/GluN2A and GluN1-4a/GluN2A receptors (Fig. 2*H*, Table 3). Finally, we found that the Po values determined for WT and mutant YFP-GluN1/GluN2A receptors did not correlate with their surface expression levels (r = 0.20, p = 0.75, R^2^ = 0.04; Fig. S5). Our results showed that reduced surface expression of GluN1/GluN2A receptors with disrupted disulfide bonds in the GluN1 subunit does not correlate with the EC_50_ values for L-glutamate or glycine, nor with Po values.

In a parallel approach, we studied whether disruption of disulfide bonds in the GluN1 subunit alters the surface expression and functional properties of the GluN1/GluN2B receptors. Using immunofluorescent labeling with an anti-GFP antibody, we observed that WT and mutant YFP-GluN1-1a/GluN2B receptors exhibited the following order of surface expression levels: WT YFP-GluN1-1a/GluN2B > YFP-GluN1-1a-C744S-C798S/GluN2B > YFP-GluN1-1a-C79S-C308S/GluN2B > YFP-GluN1-1a-C420S-C454S/GluN2B > YFP-GluN1-1a-C436S-C455S/GluN2B (Fig. 3*A,B*), similar to the GluN1/GluN2A receptors (Fig. 2*B* and S2). Our electrophysiological experiments showed that HEK293T cells transfected with YFP-GluN1-1a-C436S-C455S/GluN2B receptors did not produce any current responses; therefore, we further examined the YFP-GluN1-4a-C436S-C455S/GluN2B receptors (Fig. 3*C-D*). The analysis of the WT and mutant YFP-GluN1/GluN2B receptors and untagged GluN1-4a/GluN2B receptors showed that i.) GluN1-C79S-C308S/GluN2B and GluN1-C436S-C455S/GluN2B receptors exhibited higher EC_50_ value for L-glutamate, while GluN1-C744S-C798S/GluN2B receptors exhibited lower EC_50_ value for L-glutamate (Table 1) and ii.) GluN1-C79S-C308S/GluN2B receptor had a higher EC_50_ value for glycine (Table 2) when compared to WT GluN1/GluN2B receptors. In addition, we showed that the presence of the YFP tag did not affect the EC_50_ values for L-glutamate (Table 1) or glycine (Table 2) measured at the WT and mutant GluN1/GluN2B receptors. Using linear regression, we observed no statistically significant correlation of surface expression levels with EC_50_ values for L-glutamate (r = -0.57, p = 0.32, R^2^ = 0.32; Fig. S6) or glycine (r = -0.27, p = 0.66, R^2^ = 0.07; Fig. S7) for WT and mutant YFP-GluN1/GluN2B receptors. Our electrophysiological measurements with 1 µM MK-801 revealed that i.) GluN1-C79S-C308S/GluN2B receptors exhibited higher τ_w-MK-801_ values, while GluN1-C744S-C798S/GluN2B receptors exhibited lower τ_w-MK-801_ values compared to WT GluN1/GluN2B receptors (Fig. 3*E*; Table 3). Consistent with our findings with the GluN1/GluN2A receptor, the presence of the YFP tag did not affect the τ_w-MK-801_ values measured for WT and mutant GluN1/GluN2B receptors (Table 3). Finally, we observed that the calculated Po values of WT and mutant YFP-GluN1/GluN2B receptors did not correlate with their surface expression levels (r = 0.09, p = 0.88, R^2^ = 0.01; Fig. S8). Our experiments showed that reduced surface expression of GluN1/GluN2B receptors with disrupted disulfide bonds in the GluN1 subunit did not correlate with their EC_50_ values for L-glutamate or glycine, nor with Po values.

**Figure 3.**
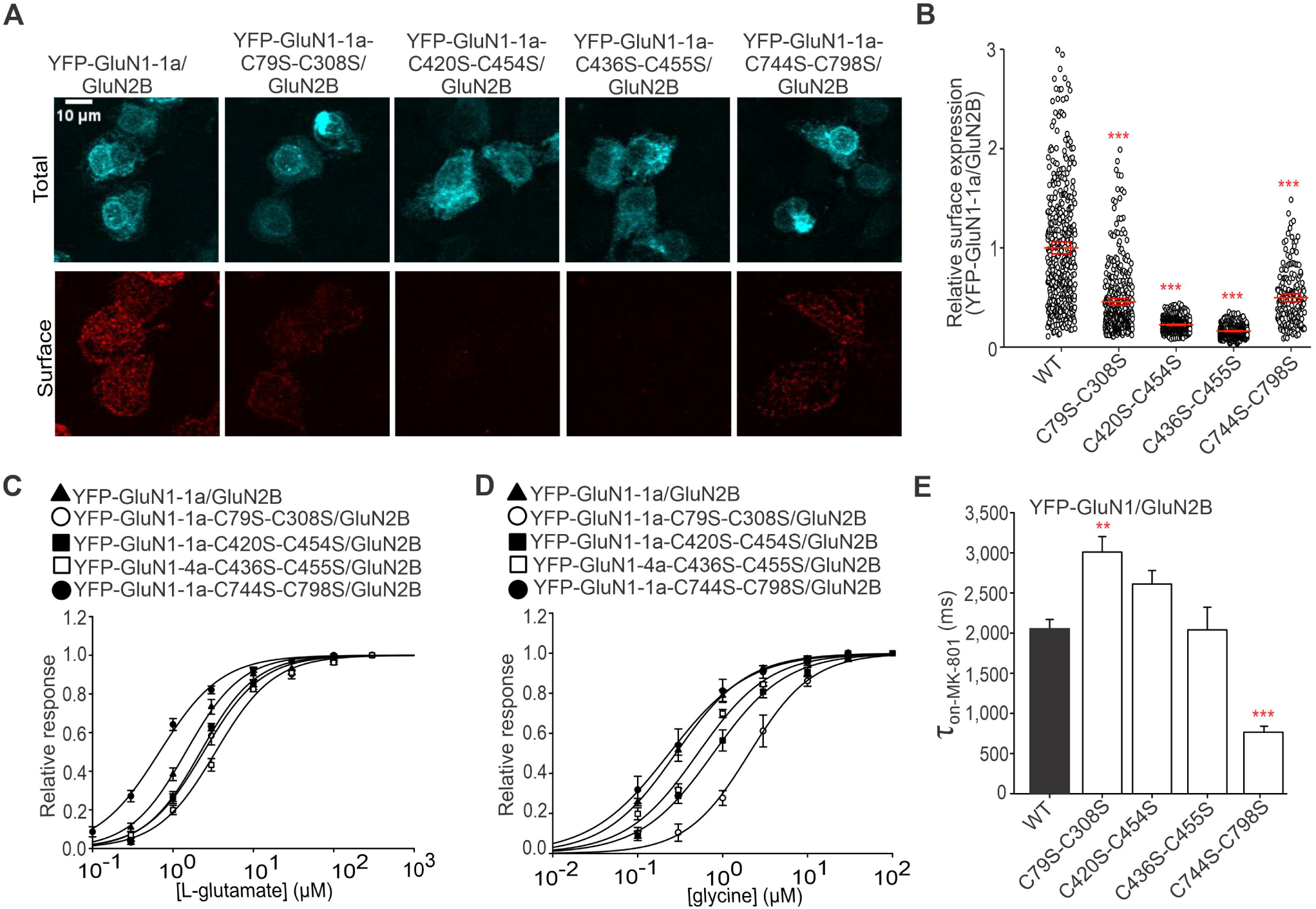
Mutations of cysteine residues forming disulfide bonds within the GluN1 subunit affect surface expression and functional properties of GluN1/GluN2B receptors. ***A,*** Representative images of HEK293T cells co-expressing the WT or mutated YFP-GluN1-1a subunit and the GluN2B subunit. The total and the cell surface number of YFP-GluN1-1a subunits (top and bottom row, respectively) were labeled using an anti-GFP antibody 24 hours after the transfection. ***B,*** Summary of the relative surface expression of NMDARs consisting of WT or mutated YFP-GluN1-1a subunit co-expressed with the GluN2B subunit measured using fluorescence microscopy; one-way ANOVA, *F*(4, 1790) = 596.75, *p* < 0.001; post hoc Tukey’s tests: *\*\*\*p* < 0.001 for WT vs. mutated NMDARs. Data points represent individual cells (*n* ≥ 184), and the red box plot represents mean ± SEM. ***C, D,*** Normalized concentration-response curves for L-glutamate (C) and glycine (D) measured from HEK293T cells expressing NMDARs containing WT or mutated YFP-GluN1-1a (YFP-GluN1-4a) subunit together with the GluN2B subunit. The data were fitted using *Equation 1* (see Methods); for a summary of fitting parameters, see Tables 1 and 2; for correlations with relative surface expression, see Figures S6 and S7, respectively. ***E,*** Summary of the τ_on-MK-801_ values for NMDARs consisting of WT or mutated YFP-GluN1-1a subunit co-expressed with the GluN2B subunit obtained by fitting the data using *Equation 2* (see Methods, Table 3); one-way ANOVA, *F*(4, 21) = 28.80, *p <* 0.001; post hoc Tukey’s tests: *\*\*p* < 0.010, *\*\*\**p < 0.001 for WT vs. mutated NMDARs. The correlation of relative surface expression and Po values (see Methods) is shown in Figure S8.

### Disruption of disulfide bonds in the GluN1 subunit reduces surface expression of unconventional GluN1/GluN3A receptors

Subsequently, we examined whether disruption of disulfide bonds in the GluN1 subunit affects the surface expression of unconventional GluN1/GluN3A receptors. We previously reported increased glycine-induced current responses of the GluN1/GluN3A receptors with the GluN1-4a splice variant (Skrenkova et al., 2018). Additionally, the GFP tag in the GluN3A subunit did not affect the τ_w-des_ values (Skrenkova et al., 2018, 2019). Therefore, we further employed WT and mutant GluN1-4a/GFP-GluN3A receptors expressed in the HEK293T cells (Fig. 4*A*). Analysis of microscopy data obtained after immunofluorescence labeling with anti-GFP antibody showed the following order of surface expression levels: WT GluN1-4a/GFP-GluN3A > GluN1-4a-C744S-C798S/GFP-GluN3A > GluN1-4a-C79S-C308/GFP-GluN3A > GluN1-4a-C420S-C454S/GFP-GluN3A > GluN1-4a-C436S-C455S/GFP-GluN3A (Fig. 4*B*). Our electrophysiological measurements showed that HEK293T cells transfected with three combinations of mutant receptors, GluN1-4a-C79S-C308/GFP-GluN3A, GluN1-4a-C420S-C454S/GFP-GluN3A, GluN1-4a-C436S-C455S/GFP-GluN3A, did not exhibit current responses >50 pA induced by glycine application in the range of 30-10,000 µM (data not shown), likely due to their low surface expression levels. To explore whether these three mutant GluN1-4a/GFP-GluN3A receptors can form functional heterotetramers, we employed 500 nM CGP-78608, a compound known to robustly potentiate the current responses of the GluN1/GluN3A receptors (Grand et al., 2018) (Fig. 4*C*). We observed that the HEK293T cells co-transfected with three mutant GluN1-4a/GFP-GluN3A receptors produced glycine-induced responses in the presence of CGP-78608, with peak current amplitudes arranged in the following order: WT GluN1-4a/GFP-GluN3A > GluN1-4a-C79S-C308/GFP-GluN3A > GluN1-4a-C420S-C454S/GFP-GluN3A > GluN1-4a-C436S-C455S/GFP-GluN3A (Fig. 4*D*). For GluN1-4a-C744S-C798S/GFP-GluN3A receptors, detectable current responses were observed after rapidly applying 10-10,000 µM glycine (Fig. 4*E*). Analysis revealed an increase in EC_50_ but not τ_w-des_ values for GluN1-C744S-C798S/GFP-GluN3A receptors compared to WT GluN1-4a/GFP-GluN3A receptors (Fig. 4*F,G*; summarized in Table 4). In conclusion, we showed that disruption of disulfide bonds in the GluN1 subunit differently reduced the surface expression of unconventional GluN1/GluN3A receptors.

**Figure 4.**
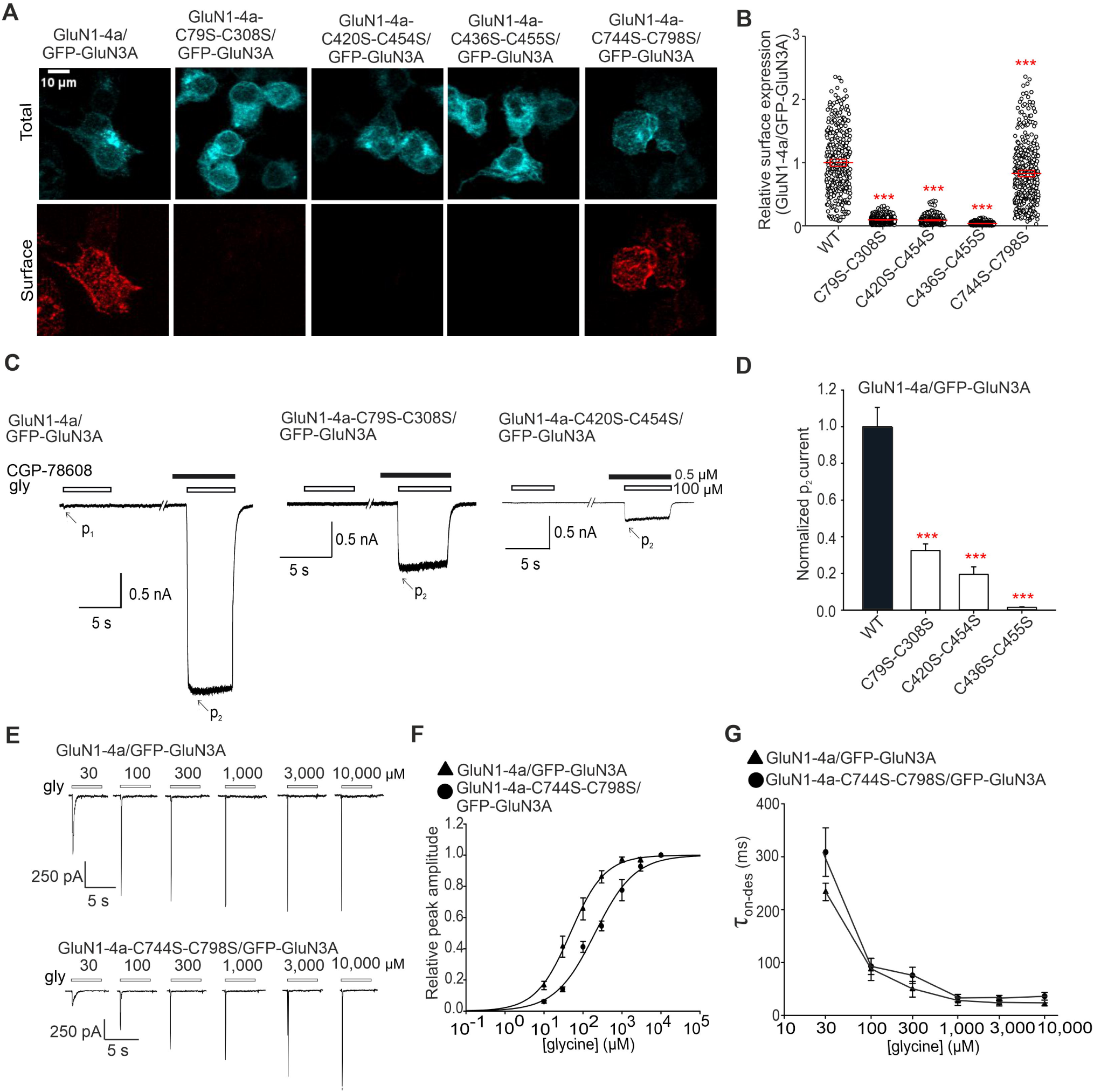
Mutations of cysteine residues forming disulfide bonds within the GluN1 subunit affect surface expression and functional properties of unconventional NMDARs. ***A,*** Representative images of HEK293T cells co-expressing the WT or mutated GluN1-4a subunit and the GFP-GluN3A subunit. The total and the cell surface number of the GFP-GluN3A subunits were labeled using an anti-GFP antibody 24 hours after the transfection. ***B,*** Summary of the relative surface expression of NMDARs consisting of WT or mutated GluN1-4a subunit co-expressed with the GFP-GluN3A subunit, measured using fluorescence microscopy; one-way ANOVA, *F*(4, 1720) = 1675.24, *p* < 0.001; post hoc Tukey’s tests: *\*\*\**p < 0.001 for WT vs. mutated NMDARs. Data points represent individual cells (*n* ≥ 391), and the red box plot represents mean ± SEM. ***C,*** Representative whole-cell voltage-clamp recordings of HEK293T cells expressing the indicated GluN1-4a/GFP-GluN3A receptors showing responses unmasked by CGP-78608. 0.5 µM CGP-78608 (black bar) was pre-applied and co-applied with the 100 µM glycine (gly; empty bar). The mark p_1_ indicates the peak current response in glycine alone, while p_2_ indicates the peak current response in the presence of both glycine and CGP-78608. ***D,*** Summary of the peak current values (nA) in response to 100 µM glycine measured in the presence of 0.5 µM CGP-78608 (p_2_) for non-conventional NMDARs consisting of either WT or mutated GluN1-4a subunit together with the GFP-GluN3A normalized to WT p_2_ peak current response (n ≥ 5); one-way ANOVA, *F*(3, 17) = 53.50, *p* < 0.001; post hoc Tukey’s tests *\*\*\*p* < 0.001 for WT vs. mutated NMDARs. ***E,*** Representative whole-cell patch-clamp recordings of HEK293T cells expressing the indicated GluN1-4a/GFP-GluN3A receptors. ***F,*** Normalized concentration-response curves for glycine obtained from HEK293T cells expressing the indicated NMDAR subunits. The data were fitted using *Equation 1* (see Methods); for a summary of fitting parameters, see Table 4. ***G,*** Summary of the τ_w-des_ measured in response to glycine in HEK293T cells expressing the indicated NMDAR subunits.

**Table 4.** Summary of the glycine concentration-response relationship analysis at WT and mutated recombinant unconventional NMDARs expressed in HEK293T cells.

| Receptor | glycine EC<sub>50</sub><br>(μM) | *h* | *n* |
| --- | --- | --- | --- |
| YFP-GluN1-1a/<br>GluN3A | 52.31 ± 12.95 | 1.26 ± 0.15 | 4 |
| YFP-GluN1-1a-C744S-C798S/<br>GluN3A | 213.41 ±<br>30.76** | 0.88 ± 0.09 | 4 |

Data are presented as mean ± SEM; values of EC<sub>50</sub> (in μM) and Hill coefficients (*h*) were obtained by fitting the data using Equation 2; (n) corresponds to the number of cells analyzed; $t_6 = -4.83$ , \*\* $p = 0.003$ (Student's *t*-test).

### Synchronized release from the ER demonstrates the importance of disulfide bonds in the GluN1 subunit in early trafficking of NMDA receptors

To determine whether the observed changes in surface expression of conventional and unconventional NMDARs with disrupted disulfide bonds are related to changes in their early transport, we performed experiments using the ARIAD system, which allows synchronized release from the ER after adding the AL (Fig. 5*A*) (Rivera et al., 2000; Hangen et al., 2018). We co-expressed HEK293T cells with WT and mutant ARIAD-GluN1-1a constructs lacking specific disulfide bonds and untagged WT GluN2A or WT GluN3A subunits. AL was added for 60 min, a time interval selected based on our recent data that showed that both ARIAD-GluN1-1a/GluN2A and ARIAD-GluN1-1a/GluN3A receptors reached sufficient numbers on the cell surface 60 min after AL addition (Netolicky et al., 2025). Then, we fixed the HEK293T cells by PFA and labeled the surface NMDARs with anti-mNEON antibodies (Fig. S9). Analysis of the microscopy data showed that the surface expression levels for both receptor combinations, ARIAD-GluN1-1a/GluN2A (Fig. 5*B*) and ARIAD-GluN1-1a/GluN3A (Fig. 5*C*), exhibited the same trends as we observed above (Fig. 2*B* and 4*B*), specifically: WT ARIAD-GluN1-1a > ARIAD-GluN1-1a-C744S-C798S > ARIAD-GluN1-1a-C79S-C308S > ARIAD-GluN1-1a-C420S-C454S > ARIAD-GluN1-1a-C436S-C455S. These experiments indicated that the disruption of disulfide bonds in the GluN1 subunit reduces the early trafficking of GluN1/GluN2A and GluN1/GluN3A receptors.

**Figure 5.**
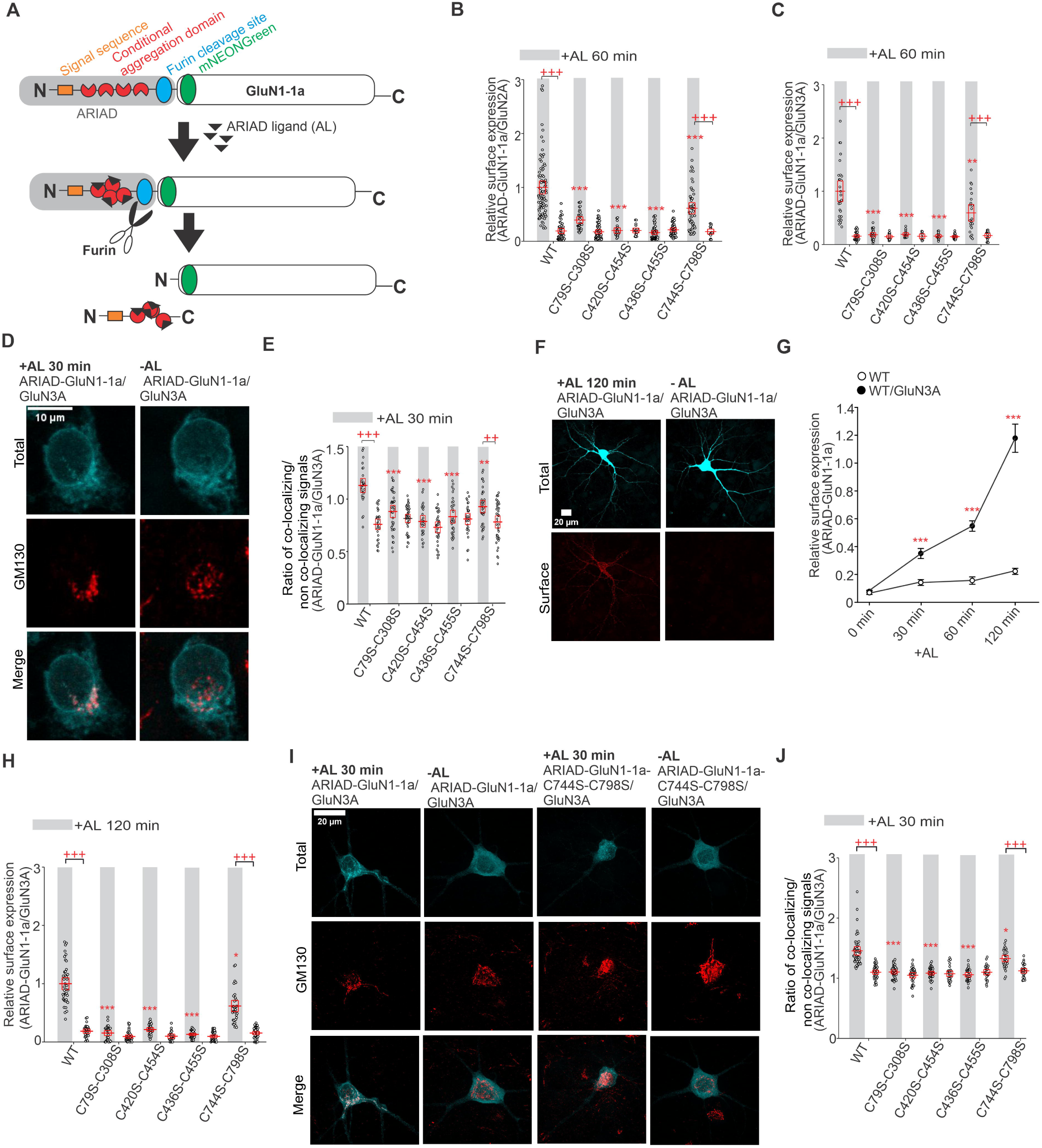
Mutations of cysteine residues forming disulfide bonds within the GluN1 subunit affect the early trafficking of conventional and unconventional NMDARs. ***A,*** Schematic representation of an ARIAD-GluN1-1a construct with a signal sequence, conditional aggregation domain, furin cleavage site, mNEONGreen (mNEON sequence was inserted after the 21st amino acid residue of the GluN1-1a), and GluN1-1a subunit. Upon the addition of the ARIAD ligand (AL) to the cells, AL binds to the conditional aggregation domain, leading to a conformational change and release of the ARIAD-mNEON-GluN1-1a construct from the ER, followed by cleavage of the ARIAD sequence by the protease furin. See also the Materials and Methods section. ***B, C,*** Summary of the relative surface expression of NMDARs consisting of WT or mutated ARIAD-GluN1-1a subunit co-expressed with the GluN2A subunit (B) or WT or mutated ARIAD-GluN1-1a subunit co-expressed with the GluN3A subunit (C), measured using fluorescence microscopy. The grey line represents 60 min in the presence of AL. For ARIAD-GluN1-1a/GluN2A: two-way ANOVA revealed a significant effect of mutation (*F*(4, 420) = 17.76, *p* < 0.001) and a significant effect of AL (*F*(1, 420) = 86.81, *p* < 0.001); for ARIAD-GluN1-1a/GluN3A: two-way ANOVA revealed a significant effect of mutation (*F*(4, 192) = 23.74, *p <* 0.001) and a significant effect of AL (*F*(1, 192) = 72.32, *p <* 0.001); post hoc Tukey’s tests: *\*\*p* < 0.010 and *\*\*\*p* < 0.001 for differences between WT and mutated ARIAD-GluN1-1a/GluN2A or ARIAD-GluN1-1a/GluN3A receptors in the presence of AL; *+++p* < 0.001 for differences between the absence and presence of AL. Data points represent individual cells (n ≥ 10), and the red box shows the mean ± SEM. For representative images, see Figure S9. ***D,*** Representative microscopy images of the HEK293T cells co-expressing WT ARIAD-GluN1-1a and GluN3A subunit in the presence (30 min) or absence of AL. The total number of NMDARs labeled with the anti-NEON antibody (top), the Golgi apparatus structures labeled with the anti-GM130 antibody (middle), and the merged image (bottom) are shown. ***E,*** Summary of the average intensity of ARIAD-hGluN1-1a subunit signal co-localized with GM130 over the average intensity of ARIAD-GluN1-1a subunit signal outside the GM130 signal, calculated for the indicated NMDAR combinations. The grey line represents 30 min in the presence of AL; two-way ANOVA revealed a significant effect of mutation (*F*(4, 318) = 7.25, *p* < 0.001) and a significant effect of AL (presence vs absence*; F*(1, 318) = 42.66*, p* < 0.001); post hoc Tukey’s tests: *\*p* < 0.05, *\*\*p* < 0.01*, ***p* < 0.001 for the difference between WT and mutated ARIAD-GluN1-1a/GluN3A receptors in the presence of AL; *+++p* < 0.001 for differences between absence and presence of AL. Data points represent individual cells (n ≥ 26), and the red box shows the mean ± SEM. ***F,*** Representative images of hippocampal neurons co-expressing ARIAD-mNEON-GluN1-1a and the GluN3A subunit, shown at 120 minutes after AL addition and at 0 minutes (without AL). The total and surface signals of ARIAD-GluN1-1a subunits (top and bottom row, respectively) were labeled using an anti-mNEON antibody 24 hours after the transfection. ***G,*** Summary of relative surface expression of NMDARs containing WT ARIAD-GluN1-1a subunits, with or without the GluN3A subunit, in hippocampal neurons at the specified time points; mean+SEM were calculated from individual segments (n ≥ 24). At 0 min, no difference was observed between ARIAD-GluN1-1a and ARIAD-GluN1-1a/GluN3A (*t_52_* = –0.64*, p* = 0.52), whereas ARIAD-GluN1-1a/GluN3A showed significantly higher values at 30 min (*t*CC = –4.30*, p* < 0.001), 60 min (*t*CC = –7.80*, ***p* <0.001) and 120 min (*t*CC = –8.59*, ***p* < 0.001, Student’s t-test). ***H,*** Summary of the relative surface expression of NMDARs consisting of WT or mutated ARIAD-GluN1-1a subunit co-expressed with GluN3A subunit measured using fluorescence microscopy. The grey line represents 120 min in the presence of the AL. Two-way ANOVA revealed a significant effect of mutation (*F*(4, 307) = 65.17, *p* < 0.001) and a significant effect of AL (*F*(1, 307) = 149.38, *p* < 0.001); post hoc Tukey’s tests: \**p* < 0.05 and \*\**p* < 0.001 for the difference between WT and mutated ARIAD-GluN1-1a/GluN3A receptors in the presence of AL; +++*p* < 0.001 for differences between absence and presence of AL. Data points represent individual segments (n ≥ 21), and the red box shows the mean ± SEM. ***I,*** Representative microscopy images of hippocampal neurons co-expressing WT or mutated ARIAD-GluN1-1a subunit and GluN3A subunit in the presence (30 min) or absence of AL. The total number of NMDARs labeled with the anti-NEON antibody (top), the Golgi apparatus structures labeled with the anti-GM130 antibody (middle), and the merged image (bottom) are shown. ***J,*** Summary of the average intensity of ARIAD-hGluN1-1a subunit signal co-localized with GM130 over the average intensity of ARIAD-GluN1-1a subunit signal outside the GM130 signal, calculated for the indicated NMDAR combinations. The grey line represents 30 min in the presence of the AL. Data points represent individual neurons (n ≥ 26), and the red box shows the mean ± SEM. Two-way ANOVA revealed a significant effect of mutation (*F*(4, 334) = 34.11, *p* < 0.001) and a significant effect of AL (*F*(1, 334) = 55.50, *p* < 0.001); post hoc Tukey’s tests: \**p* < 0.05, \*\*\**p* < 0.001 for the difference between WT and mutated ARIAD-GluN1-1a/GluN3A receptors in the presence of AL; +++*p* < 0.001 for differences between absence and presence of AL.

Our recent work showed that the WT ARIAD-GluN1-1a/GluN2A receptor does not co-localize with the Golgi apparatus (GA) at different times after AL addition. In contrast, the WT ARIAD-GluN1-1a/GluN3A receptor exhibits strong co-localization with the GA 30 minutes after AL addition (Netolicky et al., 2025). Thus, we further studied the rate of co-localization of WT and mutant ARIAD-GluN1-1a/GluN3A receptors with GA at time points 0 and 30 min after AL addition, using an anti-GM130 antibody on fixed HEK293T cells (Fig. 5*D*). Analysis of the microscopy data revealed the following order of co-localization rates with GA after 30 min of incubation with AL: WT ARIAD-GluN1-1a/GluN3A> ARIAD-GluN1-1a-C744S-C798S/GluN3A> ARIAD-GluN1-1a-C79S-C308S/GluN3A> ARIAD-GluN1-1a-C420S-C454S/GluN3A> ARIAD-GluN1-1a-C436S-C455S/GluN3A receptors (Fig. 5*E*). Without added AL, all WT and mutated ARIAD-GluN1-1a/GluN3A receptors showed a low co-localization rate with GA structures, at the level of the negative control (ARIAD-GluN1-1a, Fig. 5*E*). Subsequently, we examined the hippocampal neurons; co-transfection with the WT ARIAD-GluN1-1a and the WT GluN3A subunits resulted in robust surface expression of the mNEON-GluN1-1a/GluN3A receptors after AL addition (with increasing levels after 30, 60, and 120 min) but not under control conditions (0 min; Fig. 5*F,G*). Consistent with previous data showing the excess of the endogenous GluN1 subunit in the ER (Prybylowski et al., 2002), the transfection of the ARIAD-GluN1-1a construct resulted in slightly increased surface expression levels after AL addition, compared to the neurons expressing ARIAD-GluN1-1a and the WT GluN3A subunits (Fig. 5*G*). Therefore, we next compared the surface expression signals of the WT and mutant ARIAD-GluN1-1a/GluN3A receptors under control conditions (0 min) and 120 min after AL addition (Fig. 5*H*). Analysis of the microscopy data showed that the surface expression levels after AL addition followed the same trend observed in the HEK293T cells: WT ARIAD-GluN1-1a > ARIAD-GluN1-1a-C744S-C798S > ARIAD-GluN1-1a-C79S-C308S > ARIAD-GluN1-1a-C420S-C454S > ARIAD-GluN1-1a-C436S-C455S; no changes of the surface expression signals were observed among all conditions without AL addition (Fig. 5*H*). Then, we co-transfected hippocampal neurons with the WT and mutant ARIAD-GluN1-1a constructs, in combination with the WT GluN3A subunit and labeled GA with anti-GM130 antibody at 0 and 30 min after AL addition (Fig. 5*I*). Consistent with our data in HEK293T cells, our microscopy analysis showed that WT mNEON-GluN1-1a/GluN3A receptors exhibited a higher rate of co-localization with GA 30 min after addition of AL compared with control (0 min; Fig. 5*I,J*). In contrast, ARIAD-GluN1-1a-C79S-C308S/GluN3A, ARIAD-GluN1-1a-C420S-C454S/GluN3A, and ARIAD-GluN1-1a-C436S-C455S/GluN3A receptors showed similarly low rates of their co-localization with GA both 0 and 30 min after AL addition (Fig. 5*J*). Consistently, ARIAD-GluN1-1a-C744S-C798S/GluN3A receptors had ∼33% reduced co-localization signal with GA compared with WT ARIAD-GluN1-1a/GluN3A receptors 30 min after AL addition (Fig. 5*J*). These experiments support the observation that the reduction of surface expression of NMDARs with disrupted disulfide bonds occurs at the level of early trafficking, likely during their ER processing.

### Pathogenic GluN1-C744Y variant reduces surface expression of GluN1/GluN2 receptors while increasing their probability of opening

Using the database available at https://alf06.uab.es/grindb (as of February 22, 2022), we found a pathogenic variant, GluN1-C744Y (Fig. 6*A*), recently characterized by (Brock et al., 2023) and reported in ClinVar as associated with neurodevelopmental disorder with or without hyperkinetic movements and seizures. Our initial focus was on describing the functional properties of GluN1-C744Y/hGluN2A and GluN1-C744Y/hGluN2B receptors expressed in HEK293T cells through electrophysiological analysis. We found that YFP-hGluN1-1a-C744Y/hGluN2A receptors exhibited similar EC_50_ values for both L-glutamate (Fig. 6*B*) and glycine (Fig. 6*D*) compared with WT YFP-hGluN1-1a/hGluN2A receptors (Table 5). In contrast, the YFP-hGluN1-1a-C744Y/hGluN2B receptors exhibited reduced EC_50_ value for L-glutamate (Fig. 6*C*), while the EC_50_ value for glycine did not change (Fig. 6*E*), compared with WT YFP-hGluN1-1a/hGluN2B receptors (Table 5). In addition, we observed reduced τ_w-MK-801_ values for mutant YFP-hGluN1-1a-C744Y/hGluN2A and YFP-hGluN1-1a-C744Y/hGluN2B receptors compared to the corresponding WT YFP-hGluN1-1a/hGluN2A and YFP-hGluN1-1a/hGluN2B receptors (Table 6). This indicates that the pathogenic GluN1-C744Y variant increases the Po of GluN1/GluN2 receptors, which was confirmed by fitting the onset of inhibition by MK-801 to a kinetic model (Table 6). In the case of hGluN1-4a-C744Y/GFP-hGluN3A receptors, we observed detectable current responses after rapid glycine application in the 30-10,000 µM concentration range. The subsequent analysis revealed increased EC_50_ value for glycine but not τ_w-des_ value for hGluN1-4a-C744Y/GFP-hGluN3A receptors compared to WT hGluN1-4a/GFP-hGluN3A receptors (Fig. 6*F,G*; Table 5). These experiments showed that the pathogenic GluN1-C744Y variant alters the functional properties of NMDARs similarly to the NMDARs containing a double GluN1-C744S-C798S substitution.

**Figure 6.**
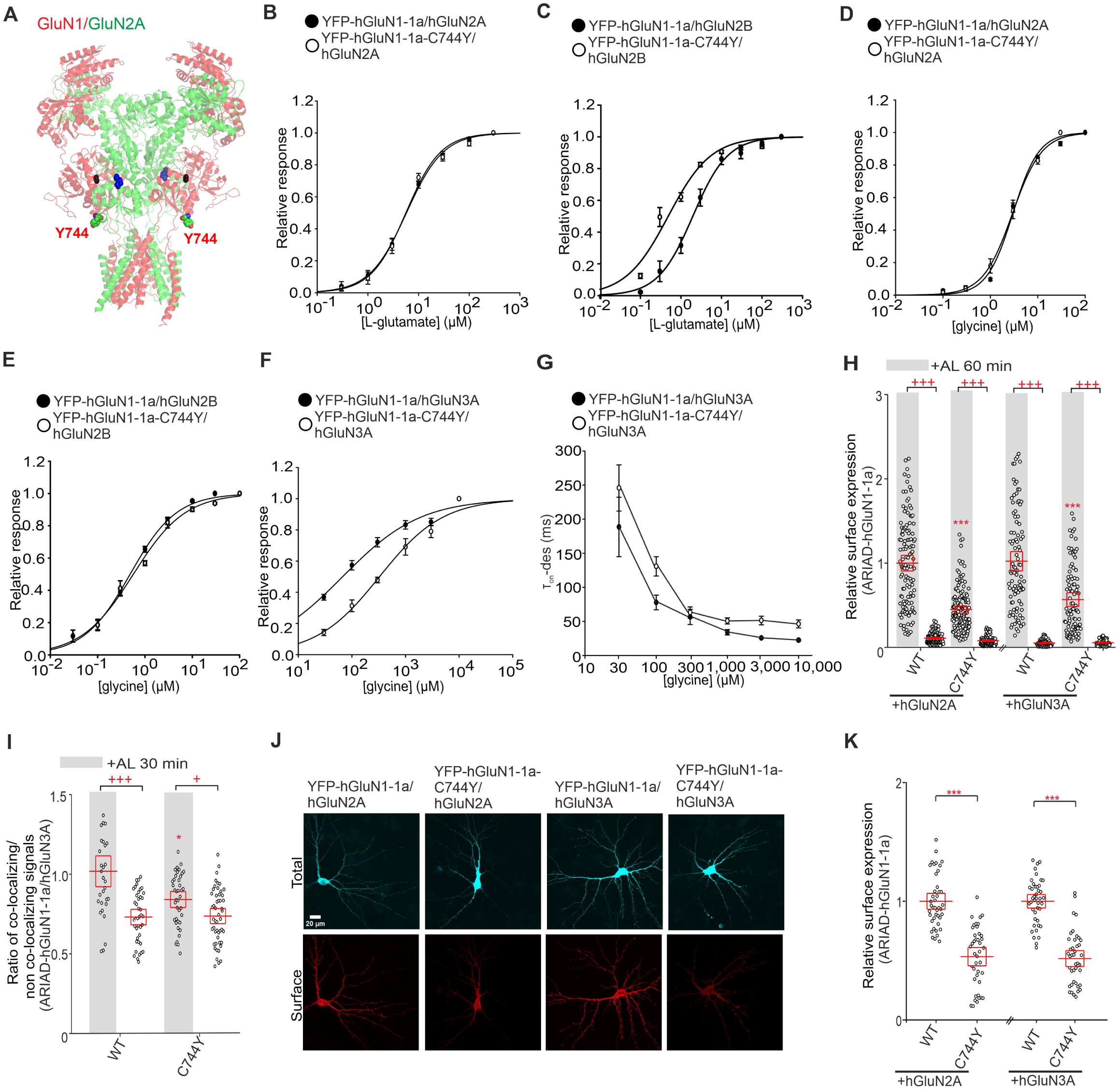
The pathogenic GluN1-C744Y variant affects the surface expression and functional properties of NMDARs similarly to the GluN1-C744S-C798S subunit. ***A,*** The structural model of the NMDAR consists of GluN1 (in red) and GluN2A (in green) subunits (PDB:7EU7) shown with L-glutamate (blue) and glycine (black) molecules bound within the LBDs and human pathogenic variant Y744. ***B, C, D, E,*** Normalized concentration-response curves for L-glutamate (B, C) and glycine (D, E) obtained from HEK293T cells expressing NMDARs containing the WT or mutated YFP-GluN1-1a subunit together with the GluN2A (B, D) or GluN2B (C, E) subunit. The data were fitted using *Equation 1* (see Methods); for a summary of fitting parameters, see Table 5. ***F,*** Normalized concentration-response curves for glycine obtained from HEK293T expressing the indicated NMDAR subunits. Data were fitted using *Equation 1* (see Methods); for a summary of fitting parameters, see Table 5. ***G,*** Summary of the τ_w-des_ measured in response to glycine in HEK293T cells expressing the indicated NMDAR subunits. ***H,*** Summary of the relative surface expression of NMDARs consisting of WT or mutant ARIAD-hGluN1-1a subunit co-expressed with hGluN2A or hGluN3A subunits in the presence (black line) or absence of AL for 60 min, normalized to the corresponding WT, measured by fluorescence microscopy. For WT and mutated ARIAD-hGluN1/hGluN2A: two-way ANOVA revealed a significant effect of mutation (*F*(1, 504) = 96.13, p < 0.001) and a significant effect of AL (*F*(1, 504) = 1221.40, p < 0.001). For WT and mutated ARIAD-hGluN1/hGluN3A: two-way ANOVA revealed a significant effect of mutation (*F*(1, 361) = 12.68, *p* < 0.001) and a significant effect of AL (*F*(1, 361) = 1164.75, p < 0.001); post hoc Tukey’s tests: \*\*\**p* < 0.001 for the differences between mutated and corresponding WT NMDAR in the presence of AL, +++*p* < 0.001 for differences between absence and presence of AL, two-way ANOVA. Data points represent individual cells (n ≥ 78), and the red box shows the mean ± SEM. The relative surface expression of the WT or mutated YFP-GluN1-1a with the WT GluN2A, WT GluN2B, and WT GluN3A subunits is shown in Figure S10. ***I,*** Summary of the average intensity of ARIAD-hGluN1-1a subunit signal co-localized with GM130 over the average intensity of ARIAD-hGluN1-1a subunit signal outside the GM130 signal, calculated for the indicated NMDAR combinations. Two-way ANOVA revealed a significant effect of mutation (*F*(1, 159) = 4.72, *p* = 0.031) and significant effect of AL (*F*(1, 159) = 35.57, *p* < 0.001); post hoc Tukey’s tests: \**p* < 0.05 for the difference between WT and mutated ARIAD-GluN1-1a/GluN3A receptors in the presence of AL; +*p* < 0.05, +++*p* < 0.001 for differences between absence and presence of AL. Data points represent individual cells (n ≥ 32), and the red box shows the mean ± SEM. ***J,*** Representative images of hippocampal neurons co-expressing the WT or mutant YFP-hGluN1-1a subunit with the hGluN2A or hGluN3A subunit. The total and the cell surface number of YFP-hGluN1-1a subunits (top and bottom row, respectively) were labeled using an anti-GFP antibody 24 hours after the transfection. ***K,*** Summary of the relative surface expression of NMDARs consisting of WT or mutated YFP-hGluN1-1a subunit co-expressed with the hGluN2A or hGluN3A subunit measured using fluorescence microscopy; *t*CC *= 8.96,* \*\*\**p* < 0.001 and *t*CC = 10.77, \*\*\**p* < 0.001; Student’s t-test. Data points represent individual segments (n ≥ 41), and the red box shows the mean ± SEM.

**Table 5.** Summary of the L-glutamate and glycine concentration-response relationship analysis at WT and mutated human recombinant NMDARs expressed in HEK293T cells.

| Receptor | L-glutamate |  |  | glycine |  |  |
| --- | --- | --- | --- | --- | --- | --- |
| | EC<sub>50</sub> (μM) | *h* | *n* | EC<sub>50</sub> (μM) | *h* | *n* |
| YFP-hGluN1-1a/<br>hGluN2A | 6.07 ± 0.29 | 1.36 ± 0.10 | 5 | 2.91 ± 0.29 | 1.46 ± 0.07 | 7 |
| YFP-hGluN1-1a-C744Y/<br>hGluN2A | 5.82 ± 0.71 | 1.33 ± 0.09 | 7 | 3.01 ± 0.41 | 1.47 ± 0.08 | 6 |
| YFP-hGluN1-1a/<br>hGluN2B | 2.24 ± 0.35 | 1.06 ± 0.06 | 6 | 0.40 ± 0.05 | 0.83 ± 0.06 | 5 |
| YFP-hGluN1-1a-C744Y/<br>hGluN2B | 0.56 ± 0.05<sup>***</sup> | 0.89 ± 0.05 | 6 | 0.59 ± 0.08 | 0.80 ± 0.03 | 5 |
| YFP-hGluN1-1a/<br>hGluN3A | - |  |  | 70.52 ± 10.66 | 0.61 ± 0.02 | 4 |
| YFP-hGluN1-1a-C744Y/<br>hGluN3A | - |  |  | 368.24 ± 51.01<sup>**</sup> | 0.75 ± 0.03 | 7 |

Data are presented as mean ± SEM; values of $EC_{50}$ (in $\mu M$ ) and Hill coefficients ( $h$ ) were obtained by fitting the data using Equation 1; ( $n$ ) corresponds to the number of cells analyzed. For WT and mutated YFP-hGluN1-1a/hGluN2B L-glutamate: $t_{10}=4.75$ , $^{**}p < 0.001$ ; for WT and mutated YFP-hGluN1-1a/hGluN3A $t_9=-4.29$ , $^{**}p = 0.002$ (Student's $t$ -test).

**Table 6.** Summary of the time constants and Po values estimated based on the inhibition onset of MK-801 at WT and mutated human recombinant NMDARs expressed in HEK293T cells.

| Receptor | $\tau_{on-MK-801}$ (ms) | $P_o$ | $n$ |
| --- | --- | --- | --- |
| YFP-hGluN1-1a/<br>hGluN2A | 789.63 ±<br>58.54 | 0.108 ±<br>0.008 | 6 |
| YFP-hGluN1-1a-C744Y/<br>hGluN2A | 240.25 ±<br>29.53<sup>***</sup> | 0.219 ±<br>0.022<sup>***</sup> | 6 |
| YFP-hGluN1-1a/<br>hGluN2B | 2008.90 ±<br>224.76 | 0.025 ±<br>0.003 | 5 |
| YFP-hGluN1-1a-C744Y/<br>hGluN2B | 846.35 ±<br>52.08<sup>***</sup> | 0.071 ±<br>0.007<sup>***</sup> | 6 |

Data are presented as mean ± SEM; values of $\tau_{w-MK-801}$ (in ms) were obtained by fitting the data using Equation 2, and values of $P_o$ were estimated based on fitting the inhibition onset of MK-801 to a kinetic model using Gepasi (see Methods; ( $n$ ) corresponds to the number of cells analyzed. For $\tau_{on-MK-801}$ : WT vs. mutated YFP-hGluN1-1a/hGluN2A $t_{10} = 8.38$ , $^{***}p < 0.001$ ; WT vs. mutated YFP-hGluN1-1a/hGluN2B $t_9 = 5.51$ , $^{***}p < 0.001$ . For $P_o$ : WT vs. mutated YFP-hGluN1-1a/hGluN2A $t_{10} = -4.74$ , $^{***}p < 0.001$ ; WT vs. mutated YFP-hGluN1-1a/hGluN2B $t_9 = -5.61$ , $^{***}p < 0.001$ (Student's $t$ -test).

To investigate the impact of the pathogenic GluN1-C744Y variant on the surface expression of NMDARs, we co-transfected the WT YFP-hGluN1-1a or mutant YFP-hGluN1-1a-C744Y subunits with the untagged hGluN2A, hGluN2B, and hGluN3A subunits in HEK293T cells and labeled them with anti-GFP antibody. Microscopy data analysis revealed that the presence of the pathogenic GluN1-C744Y variant reduced the surface expression levels of YFP-hGluN1-1a/hGluN2A, YFP-hGluN1-1a/hGluN2B and YFP-hGluN1-1a/hGluN3A receptors by ∼43%, ∼66 %, and ∼42 % (Fig. S10). To test whether the presence of the pathogenic GluN1-C744Y variant affects the early trafficking of NMDARs, we prepared ARIAD-hGluN1-1a and ARIAD-hGluN1-1a-C744Y constructs and co-expressed them with untagged hGluN2A, hGluN2B and hGluN3A subunits in HEK293T cells. Using an anti-mNEON antibody, we first verified that the ARIAD-hGluN1-1a/hGluN2A, ARIAD-hGluN1-1a-C744Y/hGluN2A, ARIAD-hGluN1-1a/hGluN3A, and ARIAD-hGluN1-1a-C744Y/hGluN3A receptor combinations did not reach the cell surface in the absence of AL (Fig. 6*H*). In contrast, 60 min after the AL addition, both ARIAD-hGluN1-1a-C744Y/hGluN2A and ARIAD-hGluN1-1a-C744Y/hGluN3A receptors exhibited ∼55% respectively ∼43% reduction in their surface expression compared to the corresponding WT ARIAD-hGluN1-1a/hGluN2A and ARIAD-hGluN1-1a/hGluN3A receptors (Fig. 6*H*). In addition, we found that the ARIAD-hGluN1-1a-C744Y/hGluN3A receptor expressed in HEK293T cells exhibited a reduced rate of co-localization with GA (labeled with anti-GM130 antibody) 30 min after AL addition compared with ARIAD-hGluN1-1a/hGluN3A receptor (Fig. 6*I*). Next, we co-transfected rat hippocampal neurons (DIV14) with WT YFP-hGluN1-1a or YFP-hGluN1-1a-C744Y subunits together with WT hGluN2A or WT hGluN3A subunits, and determined their total and surface expression levels 24 hours later by labeling with anti-GFP antibodies (Fig. 6*J*). Our analysis showed reduced surface expression of YFP-hGluN1-C744Y subunit co-expressed with the hGluN2A or hGluN3A subunits (∼53% and ∼52%, respectively) compared to the corresponding WT YFP-hGluN1 subunit (Fig. 6*K*). These experiments demonstrated that the pathogenic GluN1-C744Y variant caused a comparable reduction in the early trafficking of both conventional and unconventional NMDARs, similar to what we observed with NMDARs containing the double GluN1-C744S-C798S substitution.

We furthermore examined whether the presence of the GluN1-C744Y variant affected the extent of neuronal damage in a model of NMDA-induced excitotoxicity. We incubated infected hippocampal neurons (DIV14) expressing YFP-hGluN1-1a or YFP-hGluN1-1a-C744Y subunits for 1 hr with different concentrations of NMDA (0, 10, or 30 μM) in the presence of glycine (10 μM). We then cultured the hippocampal neurons for 23 hours and determined the proportion of dead cells by analyzing nuclear size using Hoechst 33342 (Fig. 7*A,B*). We observed a higher proportion of dead cells expressing the YFP-hGluN1-1a-C744Y subunit compared to the cells expressing the YFP-hGluN1-1a subunit after their incubation with 10 or 30 μM NMDA (Fig. 7*C* and S11). These results indicate that the presence of the GluN1-C744Y variant leads to the potentiation of neuronal death in a model of NMDA-induced excitotoxicity.

**Figure 7.**
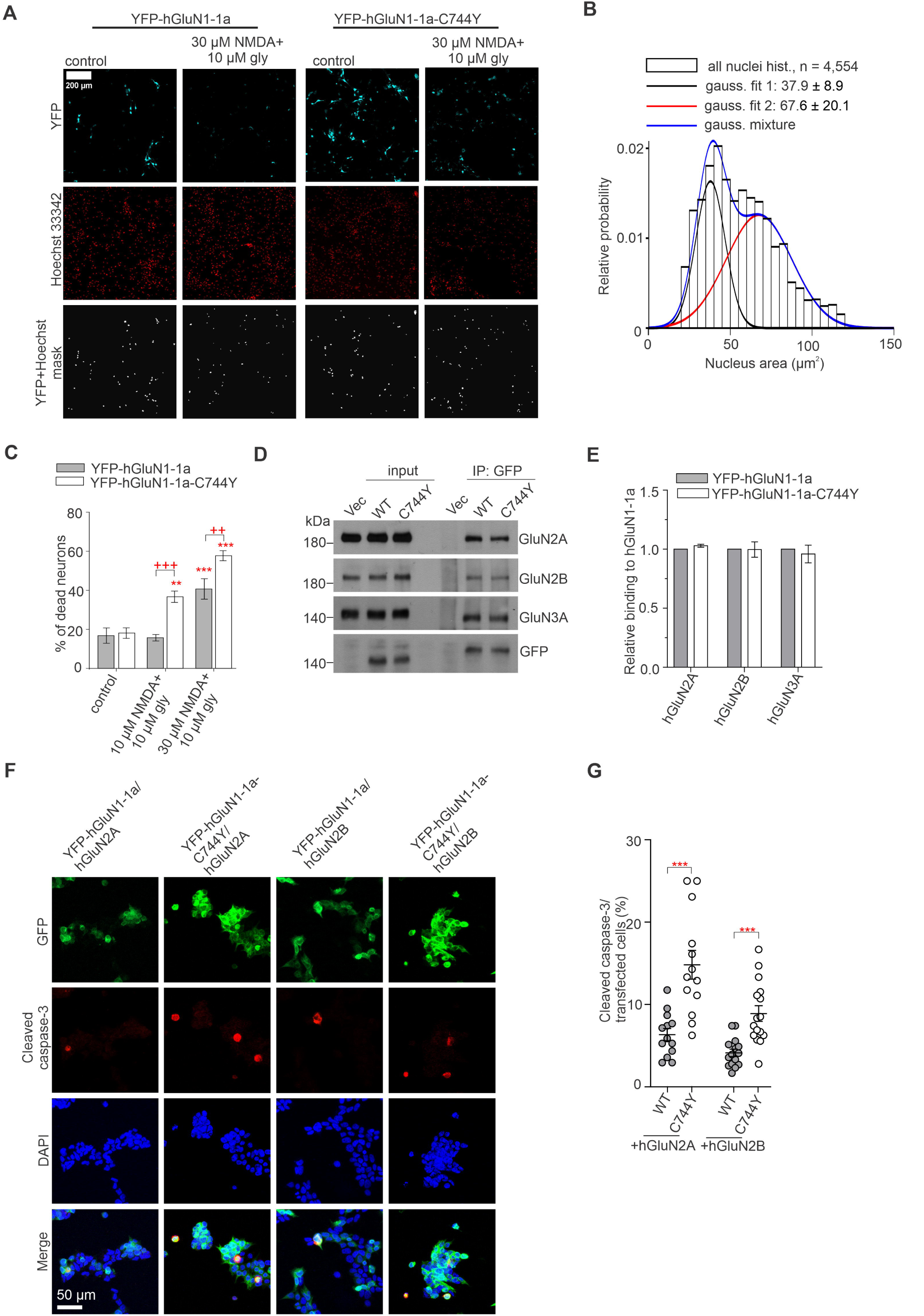
The pathogenic GluN1-C744Y variant increases NMDA-induced excitotoxicity in hippocampal neurons. ***A,*** Representative images of hippocampal neurons expressing YFP-hGluN1-1a or YFP-hGluN1-1a-C744Y treated with a control solution or 30 µM NMDA and 10 µM glycine for 1 hour. After 23 hours, YFP-positive cells were analyzed for excitotoxicity by staining with Hoechst 33342. ***B,*** Histogram of nuclear area fitted with a two-Gaussian model (blue line), separating pyknotic (black line) and non-pyknotic (red line) cell populations. For the distribution of nuclear areas, see Figure S11. ***C,*** Summary of neuronal cell death (see Methods) observed in cells expressing YFP-hGluN1-1a or YFP-hGluN1-1a-C744Y subunits after the indicated treatments (n ≥ 1,549 cells per condition collected from 2 independent experiments); two-way ANOVA revealed a significant effect of NMDA concentration (*F*(2, 114) = 45.27, *p* < 0.001) and a significant effect of mutation (*F*(1, 114) = 21.68, *p* < 0.001); post hoc Tukey’s tests: \*\**p* <0.01, \*\*\**p* <0.001 vs. corresponding control; ++*p* <0.01 +++*p* <0.001 for differences between WT and mutated YFP-hGluN1-1a. **D,** Co-immunoprecipitation assay performed in primary rat hippocampal neurons expressing WT YFP-hGluN1-1a or YFP-hGluN1-1a-C744Y subunits. Lysates were immunoprecipitated using an anti-GFP antibody, followed by western blotting with the indicated antibodies. Full-length western blot images are provided in Figure S12. **E,** Quantification of the relative binding of endogenous NMDAR subunits to WT YFP-hGluN1-1a or YFP-hGluN1-1a-C744Y subunits. Each value was normalized to the bait signal (GFP) and represents relative binding to the YFP-hGluN1-1a-C744Y subunit. Data are presented as mean ± SEM; n = 3; Student’s t-test. Note that binding of hGluN1-C744Y subunit to all subunits was reduced by approximately 15% compared to WT YFP-hGluN1-1a subunit, likely due to GluN1-C744Y-associated excitotoxicity; however, this reduction was not observed after normalization to the bait signal (GFP). **F,** Representative images of HEK293T cells co-expressing WT YFP-hGluN1-1a and YFP-hGluN1-1a-C744Y subunits and hGluN2A, hGluN2B, or hGluN3A subunits, treated with 1 mM L-glutamate and 100 µM glycine for 24 h. Excitotoxicity was assessed by cleaved caspase-3 immunostaining. **G,** Quantitation of cleaved caspase-3-positive cells per GFP-positive cells. The scatter plot presented as mean ± SEM, with each data point representing a value obtained from an image area of ∼ 200×200 mm (n ≥ 285); for WT or mutated YFP-hGluN1/hGluN2A: *t_602_* = −3.38, \*\*\**p* <0.001; for WT or mutated YFP-hGluN1/hGluN2B: *t_1108_* = −12.29, *** *p* <0.001; Student’s t-test.

We next investigated whether the GluN1-C744Y variant increases excitotoxicity by altering the subunit composition of NMDARs. The infected rat hippocampal neurons were subjected to co-immunoprecipitation and western blot analysis, which showed that the YFP-hGluN1-C744Y subunit assembled with endogenous GluN2A, GluN2B, and GluN3A subunits comparably to the WT YFP-hGluN1-1a subunit (Fig. 7*D,E*). To assess whether the GluN1-C744Y variant differently alters the excitotoxicity in the HEK293T cells expressing the hGluN1/hGluN2A or hGluN1/hGluN2B receptors, we incubated the transfected HEK293T cells with 1 mM L-glutamate and 100 µM glycine for 24 h. Cells were then stained with antibodies against GFP and activated caspase-3, as well as with DAPI to visualize nuclei (Fig. 7*F*). In both cases, we observed a higher proportion of caspase-3-positive cells expressing GluN1-1a-C744Y subunit compared to the WT hGluN1-1a subunit (Fig. 7*G*). These results indicate that the GluN1-C744Y variant increases excitotoxicity similarly at both GluN1/GluN2A and GluN1/GluN2B receptors.

It has been well established that the presence of the GluN3A subunit restricts spine maturation (Fiuza et al., 2013). We next examined whether the expression of the YFP-hGluN1-1a-C744Y subunit alone or in combination with the hGluN3A subunit affects the maturation of dendritic spines. We performed fluorescence microscopy on hippocampal neurons transfected with mCherry alone or in combination with WT YFP-hGluN1-1a, YFP-hGluN1-1a-C744Y, WT YFP-hGluN1-1a/hGluN3A, or YFP-hGluN1-1a-C744Y/hGluN3A constructs, labeled with mCherry and GFP antibodies. We classified dendritic spines as mushroom, stubby, thin, or filopodia (described in the Material and Methods section). Representative images with typical examples of each dendritic spine type are shown in Fig. 8*A*. Consistent with previous data (Fiuza et al., 2013; Kehoe et al., 2014), the co-transfection of the WT YFP-hGluN1-1a/hGluN3A receptor decreased the fraction of mushroom and stubby dendritic spines and increased the proportion of filopodia (Fig. 8*B*). By contrast, the co-transfection of YFP-hGluN1-1a-C744Y/hGluN3A receptor displayed a distribution similar to that of control conditions (mCherry, mCherry+WT YFP-hGluN1-1a, or mCherry+YFP-hGluN1-1a-C744Y), except for a slight increase of the fraction of thin dendritic spines. These data indicate that the presence of the hGluN1-1a-C744Y variant counteracts the effect of the hGluN3A subunit on decreasing dendritic spine maturation, likely by reduced surface delivery of the GluN3A-containing NMDARs in hippocampal neurons.

**Figure 8.**
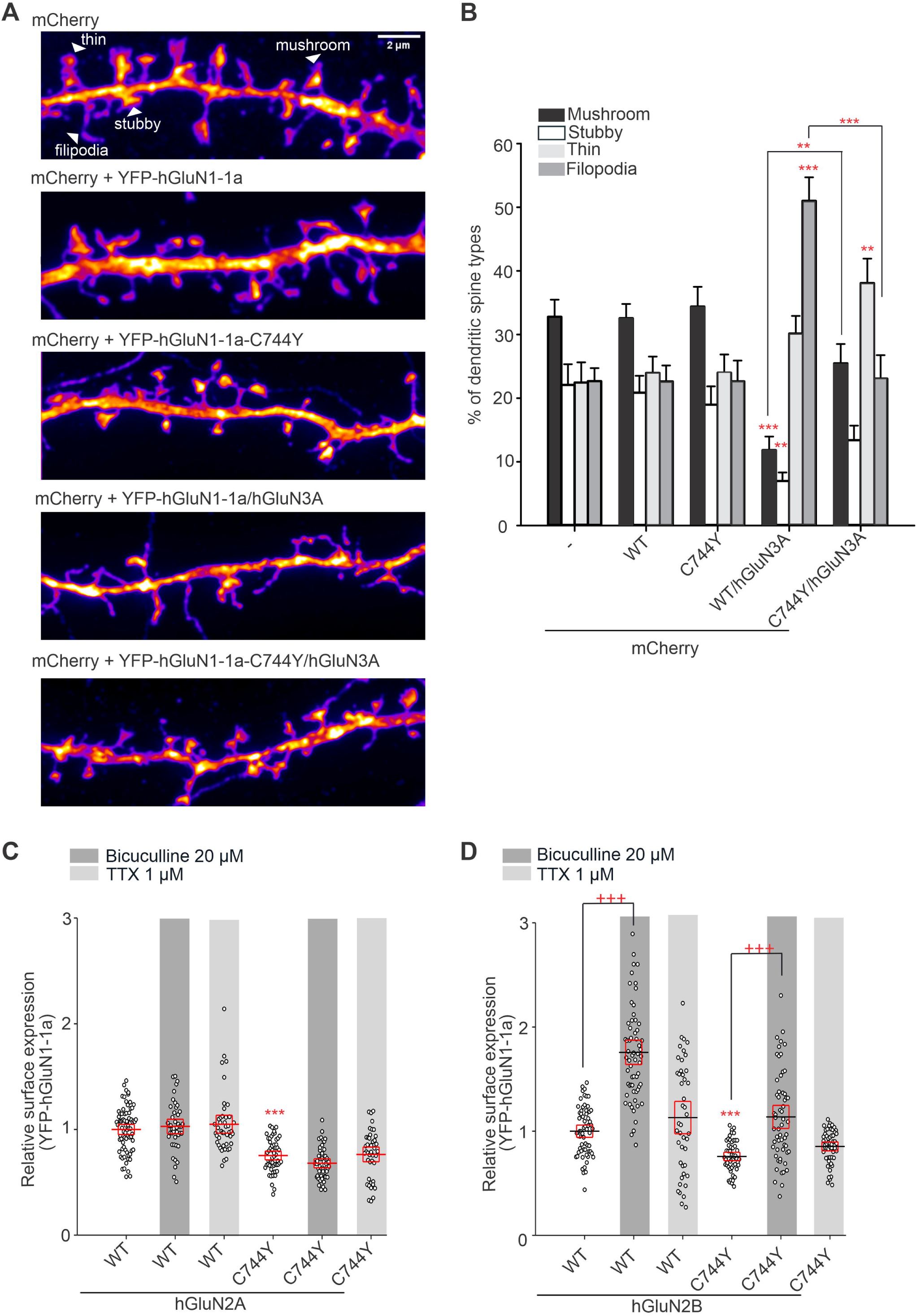
The pathogenic GluN1-C744Y variant diminishes the GluN3A subunit-induced alterations of the dendritic spine maturation. ***A,*** Representative images of hippocampal neurons expressing mCherry, and YFP-hGluN1-1a, YFP-hGluN1-1a-C744Y, YFP-hGluN1-1a/hGluN3A, and YFP-hGluN1-1a-C744Y/hGluN3A constructs with indicated types of the dendritic spines (mushroom, stubby, thin, and filopodia). **B,** Percentage distribution of dendritic spine types under the indicated conditions (n ≥ 15). One-way ANOVA for each dendritic spine type: mushroom (*F*(4,74) = 11.98, p <0.001), stubby (*F*(4,74) = 5.64, p < 0.001), thin (F(4,74) = 4.42, p = 0.003) and filopodia (*F*(4,74) = 16.28, *p* < 0.001); post hoc Tukey’s tests: \*\**p* <0.01, \*\*\**p* <0.001 vs. mCherry control or between WT and mutated YFP-hGluN1/hGluN3A. **C, D,** Relative surface expression after treatment with the indicated concentrations of bicuculline and TTX in rat hippocampal neurons expressing either YFP-hGluN1/hGluN2A (C) or YFP-hGluN1/hGluN2B (D) receptors. Two-way ANOVA for YFP-hGluN1/hGluN2A revealed a significant effect of mutation (*F*(1, 301) = 142.46, p < 0.001) and no significant effect of treatment (*F*(2,301) = 1.10, *p* = 0.335); for YFP-hGluN1/hGluN2B revealed a significant effect of mutation (*F*(1, 327) = 71.18, *p* < 0.001) and significant effect of treatment (*F*(2, 327) = 66.30, *p* < 0.001); post hoc Tukey’s tests: \*\*\**p* <0.001 vs. WT and mutated receptors without treatment, +++ *p* <0.001 vs. WT or mutated receptors with or without bicuculline treatment. Data points represent individual segments (n ≥ 42), and the red box indicates the mean ± SEM.

We further examined whether the presence of the GluN1-C744Y variant alters surface expression of the GluN2A- and GluN2B-containing NMDARs in hippocampal neurons in response to the 48-hour-long stimulation (20 μM bicuculline) or inhibition (1 μM TTX) of the synaptic activity (Fig. 8*C,D*), as employed previously (Graves et al., 2021). Consistent with our data above, we found that the YFP-hGluN1-1a-C744Y subunit exhibited lower relative surface expression compared to the WT YFP-hGluN1-1a subunit when co-expressed either with hGluN2A or hGluN2B subunits. In addition, we observed no alterations in the relative surface expression levels of the YFP-hGluN1-1a or YFP-hGluN1-1a-C744Y subunits co-expressed with the hGluN2A subunit in the presence of bicuculline or TTX. In contrast, the treatment with bicuculline, but not TTX, increased relative surface expression of both WT YFP-hGluN1-1a and YFP-hGluN1-1a-C744Y subunits co-expressed with hGluN2B subunit. These results showed that increasing synaptic activity with bicuculline promotes increased surface localization of NMDARs in a subunit-dependent manner, independently of the presence of the GluN1-C744Y variant.

Memantine, a widely used pharmacological open-channel blocker of NMDARs, is approved for treating Alzheimer’s disease and has also been tested for other diseases associated with NMDARs (Rogawski and Wenk, 2003; Robinson and Keating, 2006). Next, we measured the inhibitory concentration-response curves for memantine at WT hGluN1-4a/hGluN2A and hGluN1-4a/hGluN2B receptors and those containing the GluN1-C744Y variant at a membrane potential of -60 mV (Fig. 9*A*). The analysis revealed that the presence of the GluN1-C744Y variant decreased the IC_50_ values for memantine at both hGluN1-4a/hGluN2A (Fig. 9*B*, Table 7) and hGluN1-4a/hGluN2B receptors (Fig. 9*C*, Table 7). We also examined onset and offset kinetic parameters (τ_on_ and τ_off_ values) of the inhibitory effect mediated by 1 µM memantine at the hGluN1-4a/hGluN2A and hGluN1-4a/hGluN2B receptors containing the GluN1-C744Y variant (Fig. 9*D*). Our measurements showed that the τ_on_ values were faster while the τ_off_ values were not altered at both hGluN1-4a-C744Y/hGluN2A and hGluN1-4a-C744Y/hGluN2B receptors compared with WT hGluN1-4a/hGluN2A and hGluN1-4a/hGluN2B receptors (Fig. 9*E-F*, Table 7). These data showed that memantine is a potent antagonist of the GluN1/GluN2A and GluN1/GluN2B receptors containing the pathogenic GluN1-C744Y variant. This conclusion was further supported by an excitotoxicity assay in hippocampal neurons, where we showed that 10 µM memantine reduced neuronal death induced by 30 µM NMDA and 10 µM glycine (Fig. 9*G*).

**Figure 9.**
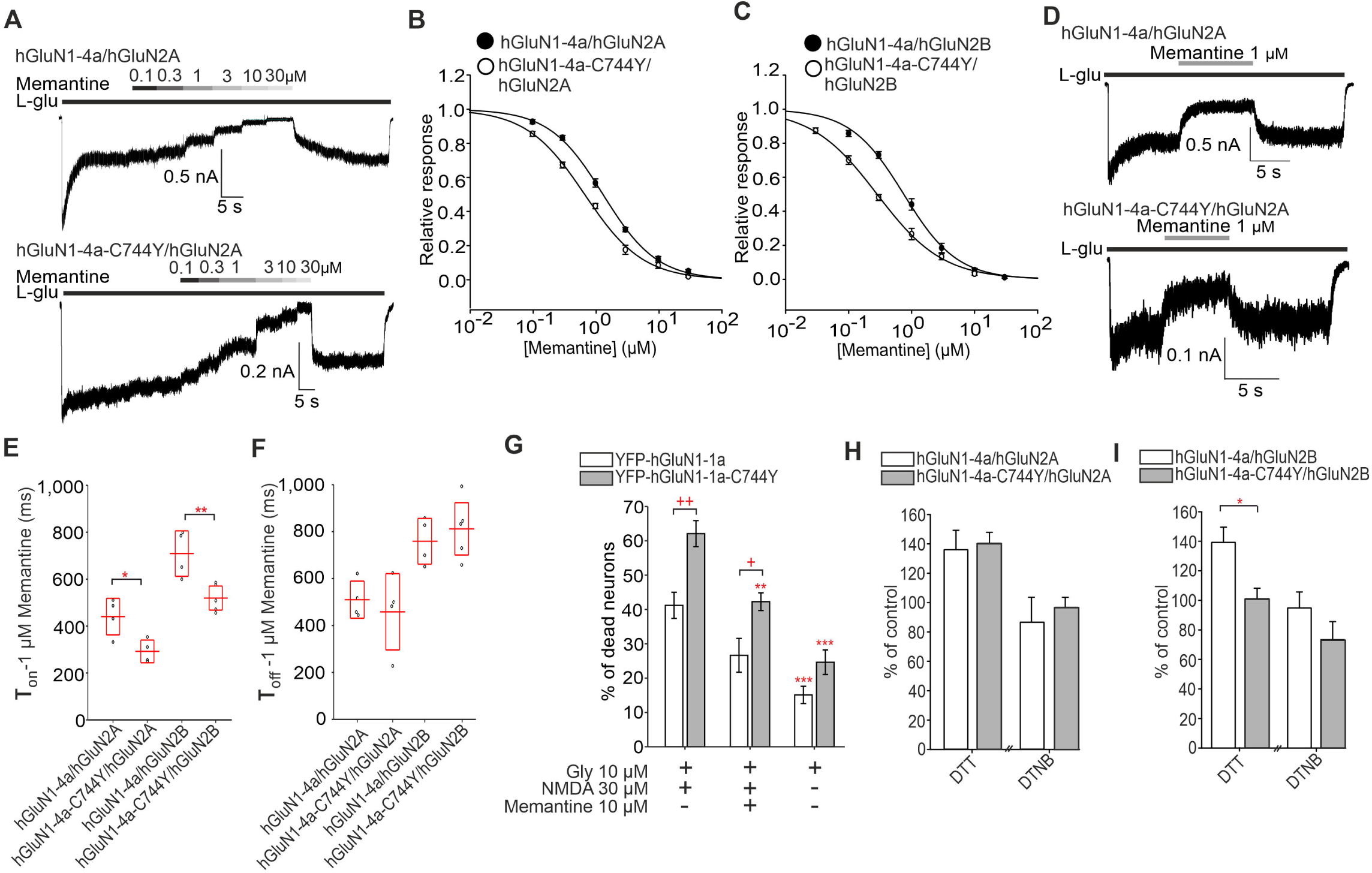
The pathogenic GluN1-C744Y variant alters memantine’s IC_50_ values and inhibitory kinetics at GluN1/GluN2 receptors. ***A,*** Representative whole-cell voltage-clamp recordings showing the concentration-response relationship of memantine in HEK293T cells expressing the indicated NMDARs. Current responses were elicited by 1 mM L-glutamate (L-glu) and 100 µM glycine (gly) and inhibited by the specified concentrations of memantine. ***B, C,*** Normalized concentration-response curves for memantine obtained from HEK293T cells co-expressing WT or mutated YFP-hGluN1-1a subunits and the hGluN2A (B) or hGluN2B (C) subunits. The data were fitted using *Equation 8* (see Methods); for a summary of fitting parameters, see Table **7**. ***D,*** Representative whole-cell voltage-clamp recordings of HEK293T cells expressing the indicated NMDAR subunits, showing the inhibition kinetics of memantine blockade. 1 µM memantine was applied in the continuous presence of 100 µM glycine (gly) and 1 mM glutamate (L-glu). ***E, F,*** Summary of the time constants for the onset (τ_on_; E) and offset (τ_off_; F) of 1 µM memantine inhibition obtained by fitting the experimental data with *Equation 8* (see Methods); fitting parameters and statistics are summarized in Table 7. ***G,*** Summary of neuronal cell death in cells expressing YFP-hGluN1-1a or YFP-hGluN1-1a-C744Y subunits after the indicated treatments (n ≥ 1,353 cells per condition collected from 2 independent experiments). Two-way ANOVA revealed a significant main effect of mutation (*F*(1,116) = 28.52, *p* < 0.001) and a significant effect of condition (*F*(2,116) = 41.74, p < 0.001); post hoc Tukey’s tests: \*\**p* < 0.01, \*\*\**p* < 0.001 vs. the corresponding control; +*p* < 0.05,++*p* < 0.01 for comparisons between WT and mutated YFP-hGluN1-1a. **H,I,** Summary of steady-state current amplitudes from HEK293T cells expressing indicated YFP-hGluN1-1a/hGluN2A (H) or YFP-hGluN1-1a/hGluN2B (I) receptors after 2 min of continuous application of 4 mM DTT, and subsequently after 2 min of continuous application of 0.5 mM DTNB, normalized to the control current amplitudes. The current responses were induced by 1 mM L-glutamate in the continuous presence of 100 µM glycine. For WT and mutated hGluN1-4a/hGluN2B receptors after DTT treatment: Student’s t-test (*t_8_*= 3.03, \**p* = 0.016); bars represent mean ± SEM (n ≥ 4).

**Table 7.** Summary of the analysis of memantine concentration-response relationships and the time constants for the onset (T_on_) and offset (T_off_) of inhibition at WT and mutated human NMDARs expressed in HEK293T cells.

| Receptor | $IC_{50}$ ($\mu M$) | $h$ | $n$ | $\tau_{on}$ (ms)<br>1 $\mu M$<br>Memantine | $\tau_{off}$ (ms)<br>1 $\mu M$<br>Memantine | $n$ |
| --- | --- | --- | --- | --- | --- | --- |
| **hGluN1-4a/**<br>**hGluN2A** | 1.34 ± 0.08 | 1.04 ± 0.06 | 6 | 440.47 ± 39.77 | 510.08 ± 40.38 | 4 |
| **hGluN1-4a-**<br>**C744Y/**<br>**hGluN2A** | 0.70 ± 0.06*** | 0.94 ± 0.06 | 7 | 292.14 ± 24.73* | 458.37 ± 83.04 | 4 |
| **hGluN1-4a/**<br>**hGluN2B** | 0.78 ± 0.09 | 1.04 ± 0.03 | 5 | 708.99 ± 49.40 | 758.93 ± 49.66 | 4 |
| **hGluN1-4a-**<br>**C744Y/**<br>**hGluN2B** | 0.28 ± 0.02*** | 0.86 ± 0.05 | 9 | 519.40 ±<br>26.19** | 811.87 ± 56.87 | 5 |

Data are presented as mean ± SEM; values of $IC_{50}$ (in $\mu M$ ) and Hill coefficients ( $h$ ) were obtained by fitting the data using Equation 8, time constants were obtained by fitting data using Equation 2; ( $n$ ) corresponds to the number of cells analyzed. For $IC_{50}$ : WT vs mutated hGluN1-4a/hGluN2A $t_{11} = 6.51$ , \*\*\* $p < 0.001$ ; WT vs mutated hGluN1-4a/hGluN2B $t_{12} = 7.11$ , \*\*\* $p < 0.001$ . For $\tau_{on}$ : WT vs mutated hGluN1-4a/hGluN2A $t_{11} = 6.51$ , \*\*\* $p < 0.001$ ; WT vs mutated hGluN1-4a/hGluN2B $t_{12} = 7.11$ , \*\*\* $p < 0.001$ . For $\tau_{on}$ : WT vs mutated hGluN1-4a/hGluN2A $t_6 = 3.17$ , \* $p = 0.019$ , WT vs mutated hGluN1-4a/hGluN2B $t_7 = 3.61$ , \*\* $p = 0.009$ . All comparisons were performed using Student's $t$ -test. The values for the WT hGluN1-4a/hGluN2B receptor were published recently (Konecny et al., 2024).

We also evaluated τ_on_ and τ_off_ values of WT and mutant hGluN1-4a/hGluN2A (3, 6, and 9 µM L-glutamate) and hGluN1-4a/hGluN2B (1, 2, and 3 µM L-glutamate) receptors in the presence of 100 µM glycine (Fig. S13*A,D*). We observed no differences between WT hGluN1-4a/hGluN2A and hGluN1-4a-C744Y/hGluN2A receptors (Fig. S13*B,C*, Table S1), whereas the hGluN1-4a-C744Y/hGluN2B receptor displayed lower τ_on_ and higher τ_off_ values compared to the WT hGluN1-4a/hGluN2B receptor (Fig. S13*E– F*, Table S1). These kinetic changes are consistent with altered EC_50_ values for L-glutamate at the GluN1/GluN2B receptor carrying the GluN1-C744Y variant.

Finally, we evaluated the redox sensitivity of the hGluN1-C744Y/hGluN2 receptors using an established redox assay (Arden et al., 1998; Ladislav et al., 2018). The current responses from the transfected HEK293T cells were evoked by the saturating concentrations of L-glutamate and glycine and this was repeated after pre-treatment with 4 mM DTT and 0.5 mM DTNB (each for 2 min). Consistent with previous data, we found that neither condition (DTT or DTNB) altered the peak current amplitudes of the WT hGluN1-4a/hGluN2A and hGluN1-4a-C744Y/hGluN2A receptors (Fig. 9*H*). In contrast, incubation with DTT, but not with DTNB, reduced the peak current amplitudes of the WT hGluN1-4a/hGluN2B receptor; however, it did not affect the hGluN1-4a-C744Y/hGluN2B receptor (Fig. 9*I*). These results showed that the redox sensitivity of the NMDARs containing the pathogenic hGluN1-4a-C744Y variant is altered in a subtype-dependent manner.

## Discussion

The existence of four disulfide bonds in the GluN1 subunit has been proposed based on structural and functional studies (Laube et al., 1993; Choi et al., 2001; Lipton et al., 2002; Furukawa and Gouaux, 2003; Papadakis et al., 2004; Kaye et al., 2007); however, their comprehensive biochemical validation has not yet been performed. In this study, we biochemically demonstrated the presence of C79-C308, C420-C454, C436-C455, and C744-C798 disulfide bonds in the GluN1 subunit. Additionally, we showed that the GluN1 subunit contains only one free cysteine residue, C459, in its extracellular region, whereas the C22 residue is absent in the mature GluN1 subunit. While the estimated molecular weight of the GluN1 subunit is 103 kDa based on its coding nucleotide sequence, after the removal of the signal peptide (18 amino acids), the actual molecular weight of the GluN1 subunit is 97 kDa—approximately 6 kDa smaller than expected. This discrepancy suggests that the mature GluN1 protein undergoes N-terminal truncation, removing approximately 60 additional residues beyond the cleavage of the signal peptide (Köpke et al., 1993). Consistent with this, our findings indicate that the N-terminus of the GluN1 subunit, including the C22 residue, is likely truncated in the mature GluN1 protein.

Concerning the effect of the disruption of the individual disulfide bonds on the functional properties of the NMDARs, we observed that GluN1-C79S-C308S and GluN1-C436S-C455S replacements caused increased EC_50_ values for L-glutamate in the case of both GluN1/GluN2A and GluN1/GluN2B receptors. Interestingly, the GluN1-C744S-C798S replacement decreased EC_50_ value for L-glutamate at GluN1/GluN2B but not GluN1/GluN2A receptors. In addition, we found that GluN1-C79S-C308S/GluN2B and GluN1-C436S-C455S/GluN2A receptors had increased EC_50_ values for glycine. A previous study showed the EC_50_ value for glycine of 2.62 µM at GluN1-C79S-C308S/GluN2A receptors (Choi et al., 2001), consistent with our reported value of 2.56 µM. Previously published EC_50_ values for GluN1/GluN2B receptors with single serine substitutions of cysteine residues forming GluN1-GluN1-C420S-C454S and C436-C455 disulfide bonds in *Xenopus* oocytes also showed similar values to our data from the HEK293T cells: GluN1-C454S/GluN2B - L-glutamate: 1.7 µM, glycine: 0.73 µM versus our data with GluN1-C420S-C454S - L-glutamate: 2.22 µM, glycine 0.88 µM; and GluN1-C436S/GluN2B - L-glutamate: 3.0 µM, glycine: 0.72 µM; GluN1-C455S/GluN2B - L-glutamate: 3.0 µM, glycine: 0.25 µM versus our data with GluN1-C436S-C455S - L-glutamate 3.27 µM, glycine 0.62 µM (Laube et al., 1993). We also found that GluN1-C744S-C798S/GluN2A, GluN1-C79S-C308S/GluN2B, and GluN1-C744S-C798S/GluN2B receptors exhibited increased Po values; this finding for GluN1-C744S-C798S/GluN2A receptor is consistent with a previous study using a single-channel measurement with GluN1-C798S/GluN2A receptors (Talukder et al., 2011). Concerning the unconventional GluN1/GluN3A receptors, we observed that disrupting GluN1-C79-C308, GluN1-C420-C454, and GluN1-C436-C455 disulfide bonds essentially eliminated glycine-induced currents in HEK293T cells which were unmasked by the CGP-78608 application; however, this did not enable us to examine their functional properties. In the case of GluN1-C744S-C798S/GluN3A receptors, we revealed altered EC_50_ values for glycine and τ_w-des_ values, which were not analyzed in a previous study (Grand et al., 2018). Together, our data suggest the disruption of individual disulfide bonds in the GluN1 subunit induces subunit-dependent structural changes in the extracellular regions of the NMDARs, which alters the transduction process, leading to ion channel opening after the interaction of agonists with their LBDs (Hansen et al., 2018). However, describing these structural alterations in detail is out of the scope of this study.

Our microscopy data revealed that the individual disruptions of disulfide bonds in the GluN1 subunit reduced the surface numbers of GluN1/GluN2A, GluN1/GluN2B, and GluN1/GluN3A receptors in the following order: WT > GluN1-C744S-C798S > GluN1-C79S-C308S > GluN1-C420S-C454S > GluN1-C436S-C455S. Using a synchronized release of the NMDARs from the ER of the HEK293T cells, we showed that changes in the surface expression of NMDARs with disrupted disulfide bonds are caused by a reduction in their early trafficking, likely at the level of ER, which is consistent with the presence of the specific machinery for the production of the disulfide bonds within the ER (Oka and Bulleid, 2013). We further documented that the ARIAD technology, in combination with NMDARs, can also be used in hippocampal neurons, as we found that the AL addition strongly increased the surface expression of the ARIAD-GluN1-1a/GluN3A receptor compared to only slightly increased surface expression of the individually expressed ARIAD-GluN1-1a subunit. This finding aligns with the observation that the GluN1 subunit but not the GluN2 subunits are expressed in excess in the ER (Fukaya et al., 2003). We chose the WT and mutant ARIAD-GluN1-1a subunit co-transfected with the GluN3A subunit for subsequent studies as i.) WT ARIAD-GluN1-1a/GluN3A receptor co-localized with GA in the HEK293T cells, ii.) the expression of the GluN3A subunit exhibits anti-apoptotic activity in contrast to the GluN2A and/or GluN2B subunits (to minimize the potential artifacts caused by excitotoxicity) (von Engelhardt et al., 2007; Nakanishi et al., 2009), and iii.) the individual disruptions of disulfide bonds in the GluN1 subunit reduced similarly the surface numbers of all studied NMDAR subtypes. We observed a substantial degree of colocalization of the WT ARIAD-GluN1-1a/GluN3A receptors with GA in hippocampal neurons, which was diminished in an identical manner with the surface expression signals of the mutated GluN1/GluN3A receptors. This showed that the disruption of the disulfide bonds alters the early trafficking of the NMDARs, likely at the level of the ER, also in hippocampal neurons. Our detailed analyses revealed no correlation between surface expression levels and EC_50_ values for L-glutamate and glycine or Po values at GluN1/GluN2A and GluN1/GluN2B receptors. Importantly, we replicated previous findings that the GluN1/GluN2A receptor exhibits approximately twice the Po value compared to the GluN1/GluN2B receptor (Erreger et al., 2005). The similar surface expression levels of the GluN1/GluN2A and GluN1/GluN2B receptors containing the disruption of the individual disulfide bonds are not surprising, given the degree of structural homology between their extracellular regions (Zhang and Luo, 2013; Vieira et al., 2020). However, we suggest that the ER processing of all GluN1/GluN2A, GluN1/GluN2B, and GluN1/GluN3A receptors with disrupted disulfide bonds is likely regulated by a shared mechanism, although the structure of the extracellular regions of the GluN1/GluN3A receptors, as well as their functional properties, differ substantially compared with GluN1/GluN2 receptors (Michalski and Furukawa, 2024).

Our experiments with the pathogenic GluN1-C744Y variant essentially replicated our findings regarding the NMDARs with double GluN1-C744S-C798S substitutions, highlighting the critical role of the GluN1-C744-C798 disulfide bond in regulating the early trafficking and functional properties of NMDARs. A recent study reported the EC_50_ value for L-glutamate of 1.1 µM and the EC50 value for glycine of 0.7 µM for GluN1-C744Y/GluN2A receptors expressed in Xenopus oocytes, while our analysis showed the EC50 value for L-glutamate of 5.8 µM and the EC_50_ value for glycine of 3.5 µM (Brock et al., 2023). The differences in the reported EC_50_ values are relatively small, considering the micromolar range, and may result from variations in expression systems and compositions of recording solutions. In addition, using a β-lactamase assay, the authors demonstrated a reduction of ∼32% in the surface expression of GluN1-C744Y/GluN2A receptors expressed in HEK293T cells, which is consistent with our microscopy analysis, which showed a reduction of 43%. Our findings that the GluN1-C744Y variant increases excitotoxicity, but does not alter the association of the GluN1 subunit with GluN2 and GluN3A subunits, are consistent with a dominant role of the increased Po value of the YFP-GluN1-C744Y/GluN2 receptors during excitotoxicity conditions. We also revealed that the hGluN1-C744Y/hGluN2B receptor is insensitive to DTT, consistent with previous data at the GluN1-C744S-C798S/GluN2B receptor (Arden et al., 1998), underscoring the importance of these cysteine residues in the redox modulation of NMDARs. Our data with the FDA-approved memantine showed promise for the potential therapy of neurodegeneration associated with the disruption of the GluN1-C744S-C798S disulfide bond. Future studies are needed to reveal the impact of the GluN1-C744Y variant at the whole-organism level, for example, by using knock-in mice. We attempted to obtain the global knock-in mouse with the GluN1-C744Y variant, but did not get any pups carrying the correctly altered *Grin1* allele. This is consistent with the obligatory role of the GluN1 subunit in the development of the CNS (Fukaya et al., 2003). The expanding number of known pathogenic variants of the cysteine residues in the GluN subunits emphasizes the need for further research on regulating NMDARs by disulfide bonds (Swanger et al., 2016; Addis et al., 2017, 2017; Platzer et al., 2017; Franchini et al., 2020). It remains to be uncovered whether pharmacological interventions, such as those tailored to specifically target NMDARs with a disrupted disulfide bond(s), can adequately correct the associated CNS dysfunctions, including the disrupted maturation of the dendritic spines. We acknowledge that future studies should also examine whether the altered redox/oxidative environment can regulate the early trafficking of NMDARs, as previous studies showed that the redox state of the NMDARs is, for example, altered during status epilepticus (Sanchez et al., 2000; Kim et al., 2017; Jeon and Kim, 2018).

## Supporting information

Supplementary all

## Acknowledgments

Supported by project registration number LX22NPO5107 (MEYS CR): Financed by EU – Next Generation EU and by project registration number CZ.02.01.01/00/22_008/0004562 (MEYS CR). This work was also supported by the Grant Agency of Charles University (GAUK: 144323). Y.H.S. J.-m.S. and S.L. were supported by the Korea Dementia Research Project through the Korea Dementia Research Center (KDRC) (Grant No. RS-2024-00332875). We acknowledge the Microscopy Service Centre (Institute of Experimental Medicine CAS) supported by the MEYS CR (LM2023050; Czech-Bioimaging).

