## Supplementary all for "The role of disulfide bonds in the GluN1 subunit in the early trafficking and functional properties of GluN1/GluN2 and GluN1/GluN3 NMDA receptors"

**EXTENDED FIGURE LEGENDS AND TABLE**

### Figure S1

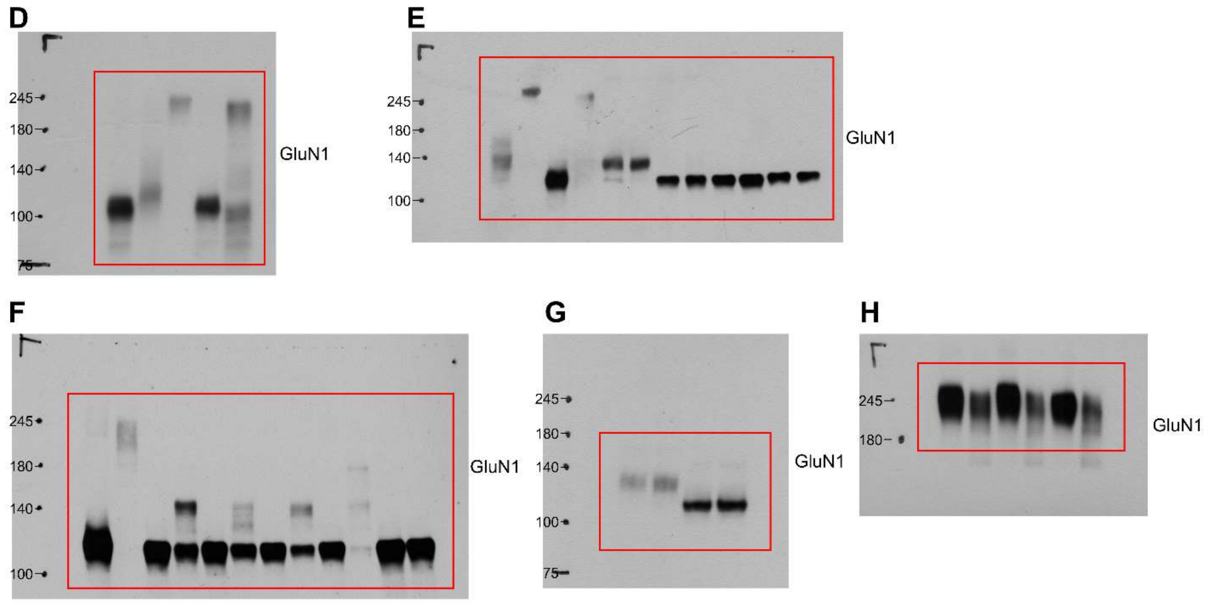


**Figure S1.** Full-length western blot images corresponding to Fig. 1. The blots shown in Fig. 1*D–H* are highlighted with red boxes.

### Figure S2

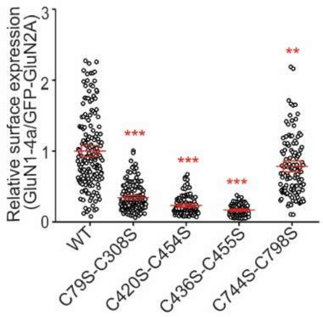


**Figure S2.** Summary of the relative surface expression of NMDARs consisting of WT or mutated GluN1-4a subunits co-expressed with the GFP-GluN2A subunit, measured using fluorescence microscopy; one-way ANOVA, $F(4, 635) = 220.81$, $p < 0.001$; post hoc Tukey's tests, $^{**}p < 0.01$, $^{***}p < 0.001$. Data points represent individual cells ($n \geq 94$), and the red box plot represents mean ± SEM.

### Figure S3

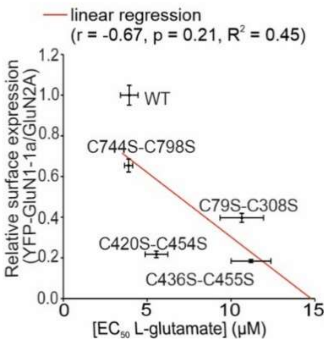


**Figure S3.** Correlation of surface expression with $EC_{50}$ values for L-glutamate in NMDARs composed of WT or mutated YFP-GluN1-1a subunit together with the GluN2A subunit. Data were fitted by linear regression ($r = -0.67$, $p = 0.21$, $R^2 = 0.45$).

### Figure S4

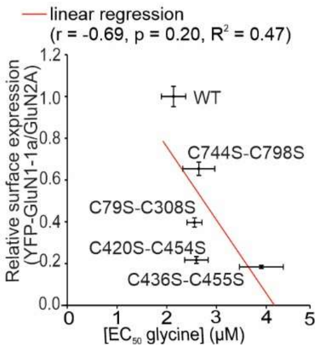


**Figure S4.** Correlation analysis of relative surface expression with $EC_{50}$ values for glycine for NMDARs composed of WT or mutated YFP-GluN1-1a subunits co-expressed with the GluN2A subunit. Data were fitted by linear regression ($r$ = -0.69, $p$ = 0.20, $R^2$ = 0.47).

### Figure S5

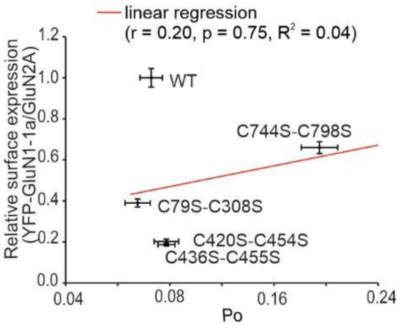


**Figure S5.** Correlation analysis of relative surface expression with the Po values determined by fitting the onset inhibition by MK-801 to the model (see the Material and Methods section, Table 3) for NMDARs containing WT or mutated YFP-GluN1-1a subunits together with GluN2A subunit; data were fitted by linear regression ($r$ = 0.20, $p$ = 0.75, $R^2$ = 0.04).

### Figure S6

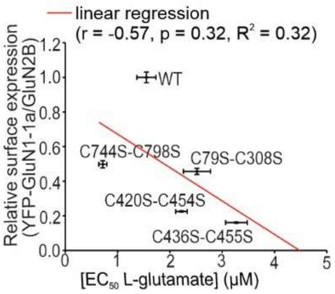


**Figure S6.** Correlation analysis of relative surface expression with $EC_{50}$ values for L-glutamate at NMDARs composed of the WT and mutated YFP-GluN1-1a subunits together with the WT GluN2B subunit. Data were fitted by linear regression (*r* = -0.57*, p* = 0.32*, $R^2$* = 0.32).

### Figure S7

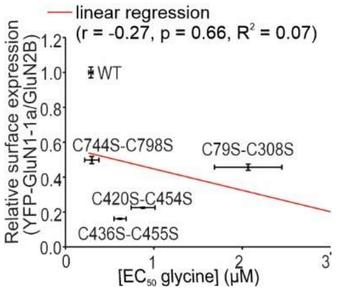


**Figure S7.** Correlation analysis of relative surface expression with $EC_{50}$ values for glycine at NMDARs composed of the WT and mutated YFP-GluN1-1a subunits together with the WT GluN2B subunit. Data were fitted by linear regression (*r* = -0.27*, p* = 0.66*, $R^2$* = 0.07).

### Figure S8

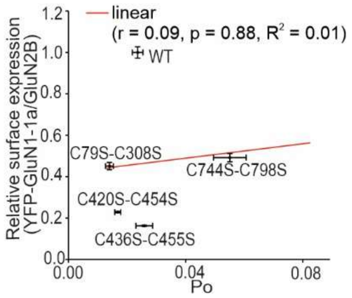


**Figure S8.** Correlation analysis of relative surface expression with the Po values determined by fitting the onset inhibition by MK-801 to the model (see the Material and Methods section, Table 3) for NMDARs containing WT or mutated YFP-GluN1-1a subunits co-expressed with the GluN2B subunit; data were fitted by linear regression (*r* = 0.09*, p* = 0.88*,* $R^2$ = 0.01).

### Figure S9

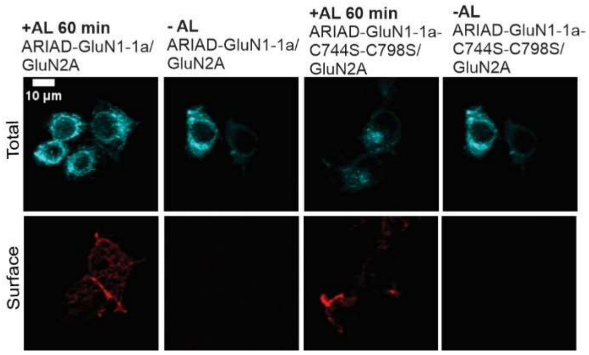


**Figure S9.** Representative images of HEK293T cells co-expressing the WT or mutated ARIAD-GluN1-1a subunit and the GluN2A subunit. The total and the surface number of the ARIAD-GluN1-1a subunits (top and bottom row, respectively) were labeled using an anti-NEON antibody 24 hours after the transfection, following 60 min incubation with or without AL.

### Figure S10

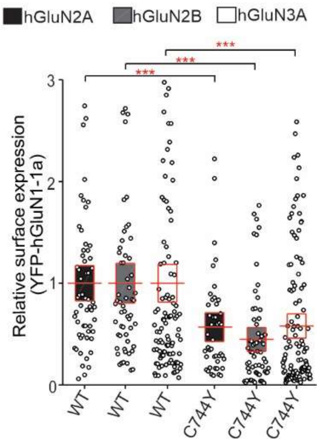


**Figure S10.** Summary of the relative surface expression of NMDARs consisting of WT or mutated YFP-hGluN1-1a subunit co-expressed with the hGluN2A, hGluN2B, or hGluN3A subunit in the HEK293T cells, normalized to the corresponding WT, as measured using fluorescence microscopy; for WT and mutated YFP-hGluN1/hGluN2A $t_{101} = 3.57$, ***$p < 0.001$, for WT and mutated YFP-hGluN1/hGluN2B $t_{117} = 4.77$, *** $p < 0.001$, and for WT and mutated YFP-hGluN1/hGluN3A $t_{233} = 3.97$, ***$p < 0.001$; Student's t-test. Data points represent individual cells ($n \geq 44$), and the red box plot shows mean ± SEM.

### Figure S11

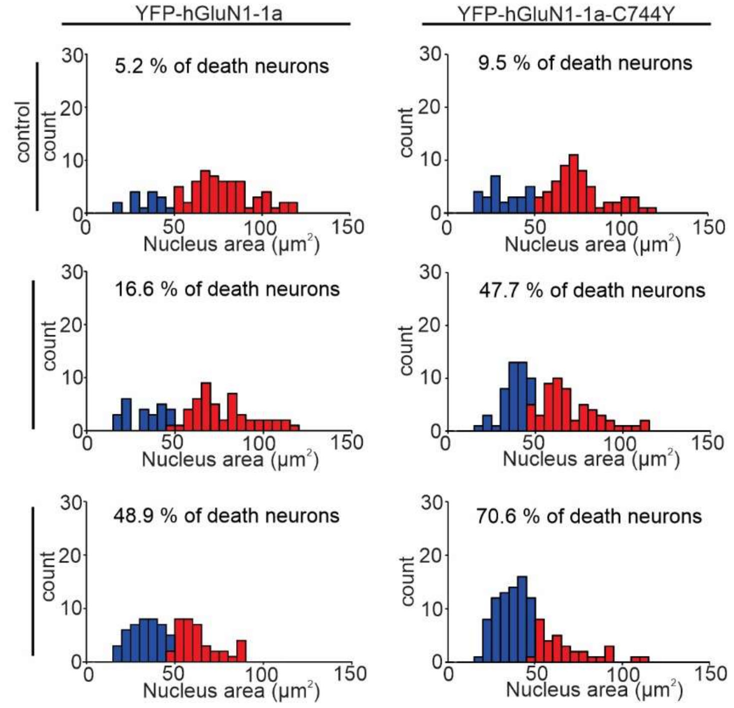


**Figure S11.** Distribution of nuclear areas in cells expressing YFP-hGluN1-1a or YFP-hGluN1-1a-C744Y subunits after the indicated treatments. Nuclei were categorized as pyknotic (blue bars) or non-pyknotic (red bars).

### Figure S12

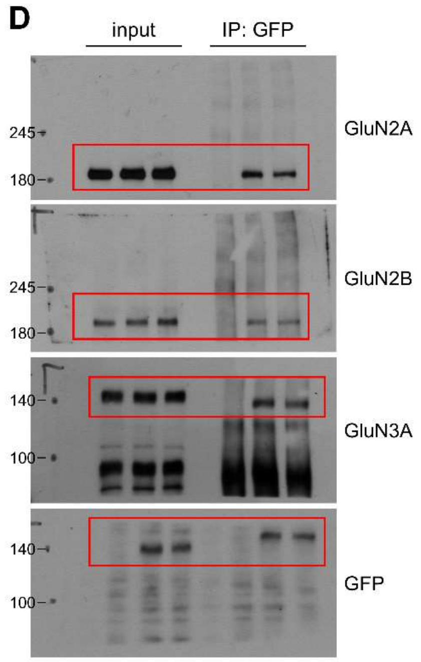


**Figure S12.** Full-length western blot images corresponding to Fig. 7. The blots shown in Fig. 7*D* are highlighted with red boxes.

### Figure S13

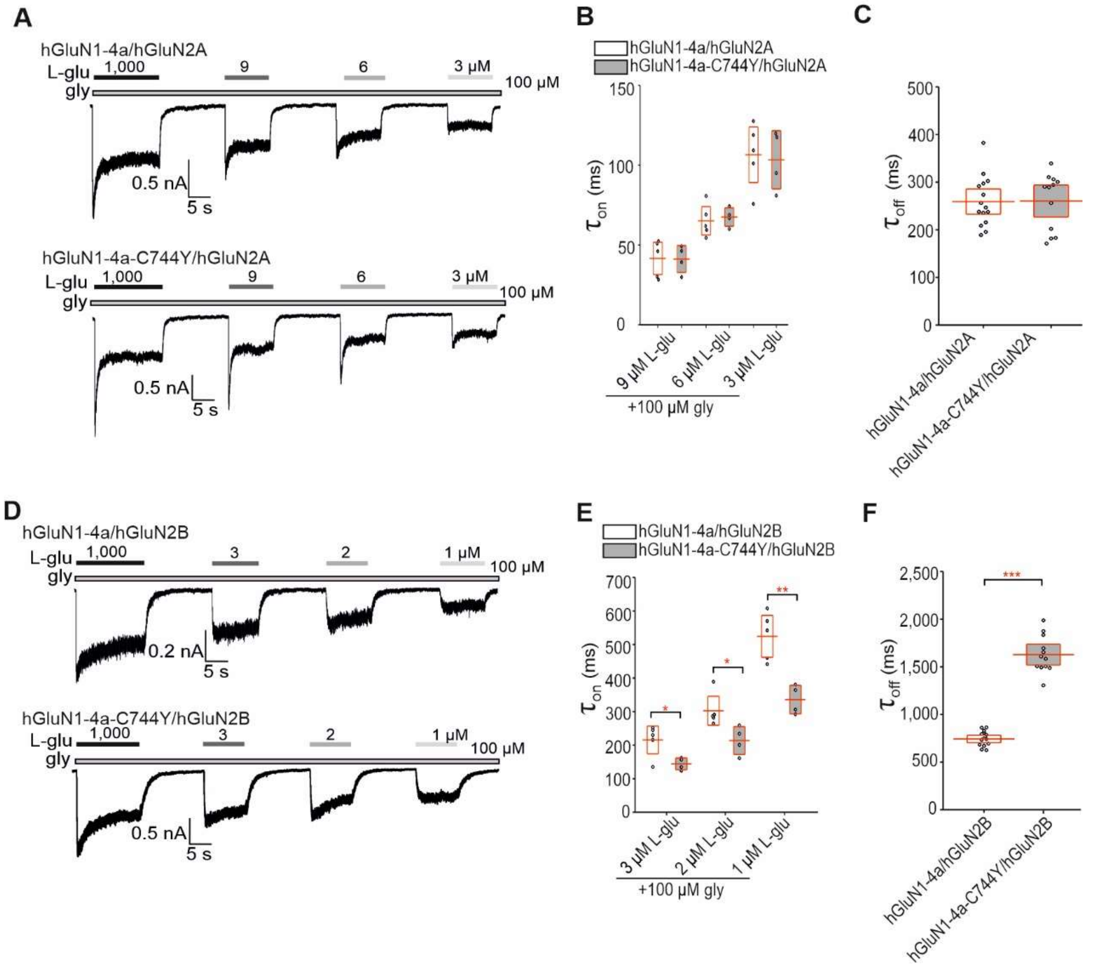


**Figure S13.** The pathogenic GluN1-C744Y variant, in combination with the GluN2B subunit, alters the kinetics of L-glutamate-evoked responses of the NMDARs.

***A,D,*** Representative whole-cell voltage-clamp recordings from HEK293T cells co-expressing WT or mutated hGluN1-4a and GluN2A (A) or GluN2B (D) subunits, showing the distinct kinetics to different concentrations of L-glutamate (L-glu) as indicated. Current responses were elicited in the continuous presence of 100 µM glycine (gly); for a summary of fitting parameters, see Tab. S1. ***B, C, E, F,*** Summary of the time constants for the onset ($\tau_{on}$; B, E) and offset ($\tau_{off}$; C, F) of the current responses induced by the indicated concentrations of L-glutamate (L-glu) obtained from HEK293T cells co-expressing the WT or mutated hGluN1-4a subunits and GluN2A (B, C) or GluN2B (E, F) subunits. The values were obtained by fitting the experimental data with *Equation 2* (see the Material and Methods); for a summary of fitting parameters and statistics, see Table S1.

**Table S1.** Summary of the time constants for the onset ($\tau_{on}$) and offset ($\tau_{off}$) of L-glutamate-induced current responses at the indicated WT and mutated human NMDARs expressed in HEK293T cells.

| Receptor | $\tau_{on}$ (ms) L-glu 9 μM | $\tau_{on}$ (ms) L-glu 6 μM | $\tau_{on}$ (ms) L-glu 3 μM | $\tau_{off}$ (ms) | n |
|---|---|---|---|---|---|
| **hGluN1-4a/ hGluN2A** | 41.62 ± 5.16 | 65.15 ± 4.17 | 106.57 ± 8.90 | 259.28 ± 20.00 | 5 |
| **hGluN1-4a-C744Y/ hGluN2A** | 41.19 ± 4.31 | 67.54 ± 2.94 | 103.38 ± 9.31 | 260.44 ± 29.39 | 4 |
| **Receptor** | **$\tau_{on}$ (ms) L-glu 3 μM** | **$\tau_{on}$ (ms) L-glu 2 μM** | **$\tau_{on}$ (ms) L-glu 1 μM** | **$\tau_{off}$ (ms)** | **n** |
| **hGluN1-4a/ hGluN2B** | 215.65 ± 21.04 | 302.60 ± 22.09 | 524.36 ± 31.90 | 742.89 ± 17.07 | 5 |
| **hGluN1-4a-C744Y/ hGluN2A** | 144.27 ± 9.02$^{*}$ | 213.79 ± 21.16$^{*}$ | 335.84 ± 21.62$^{***}$ | 1637.68 ± 87.62$^{***}$ | 4 |

*Data are presented as mean ± SEM; values of* $\tau_{on}$ *and* $\tau_{off}$ *(in ms) were obtained by fitting the data using* Equation 2*, see Methods; (n) corresponds to the number of cells analyzed. For* $\tau_{on}$ *with L-glu 3 μM in WT and mutated GluN1/GluN2B receptors (*$t_7 = 2.84$*,* $p = 0.025$*), for L-glu 2 μM (*$t_7 = 2.85$, $p = 0.025$; $t_7 = 4.62$*,* $p = 0.002$*), and for* $\tau_{off}$ *(*$t_7 = -11.28$, $p < 0.001$*); *p <0.05,* *p <0.001 for WT vs. mutated NMDAR; Student's t-te*st*.
